# mitoXplorer 3.0, a web tool for exploring mitochondrial dynamics in single-cell RNA-seq data

**DOI:** 10.1101/2024.12.17.628870

**Authors:** Margaux Haering, Andrea del Bondio, Helene Puccio, Bianca H. Habermann

**Author notes:** Corresponding author: Bianca Habermann, CNRS DR2 Address: Aix-Marseille University, CNRS, IBDM UMR7288 Parc Scientifique de Luminy 163 Avenue de Luminy 13009 Marseille, France.

## Abstract

Mitochondria are important eukaryotic organelles, best known for their function in ATP production and in cellular metabolism and signalling. It is widely accepted that their structure, composition and function differ across cell types. However, little is known about mitochondrial variability within the same cell type. To truly understand mitochondrial function and dynamics, we need to study individual cell types, as well as mitochondrial variability on a single-cell level.

Based on our mitoXplorer 2.0 web tool, we introduce mitoXplorer 3.0 with new features adapted for analysing single-cell sequencing data, focusing only on mitochondria. We provide a formatting script, scXplorer to generate mitoXplorer 3.0 compatible files for upload. This script creates pseudo-bulk transcriptomes of cell types from scRNA-seq data for differential expression analysis and subsequent mitochondria-centric analysis with mitoXplorer classical interfaces. It also creates a single-cell expression matrix only containing mitochondria-associated genes (mito-genes), which can be analysed for cell-to-cell variability with novel, interactive interfaces created for mitoXplorer 3.0: these new interfaces help to identify sub-clusters of cell types based only on mito-genes and offer in-depth mitochondria-centric analysis of subpopulations. We demonstrate the usability and predictive power of mitoXplorer 3.0 using single-cell transcriptome data from a single-cell study of Spinocerebellar Ataxia Type 1. We identified several mito-processes and mito-genes that are majorly affected in SCA1 Purkinje cells and which might contribute to our understanding of mitochondrial decline and subsequent Purkinje cell loss in this disease.

MitoXplorer 3.0 is freely available at https://mitoxplorer3.ibdm.univ-amu.fr.

## INTRODUCTION

Mitochondria are essential for eukaryotic life and are important for various cellular processes, including bioenergetics, metabolism, apoptosis, and cellular signalling [1]. While found in almost all eukaryotic cells, their structure, as well as their expression profile differs across cell types, mostly reflecting the different requirements of the cells to these organelles [2–4]. It is for this reason that particularly in highly heterogeneous tissues such as the brain, standard bulk-RNA sequencing is ill-suited to capture the expression differences of the individual cell types. Some diseases or dysfunctions also target specific cell types [5,6] and cell-type specificity is particularly important in neurodegenerative disorders like cerebellar ataxias, where disease phenotypes are often due to the perturbation or the loss of a single cell type [7]. Moreover, mitochondria are known to exhibit heterogeneity even within seemingly homogeneous cell populations [8] and such cellular heterogeneity can influence disease progression and treatment response.

With the advent of single-cell (scRNA-seq) / single-nuclei RNA sequencing (snRNA-seq), it became possible to apprehend cell-type specific expression patterns in complex tissues and to uncover cellular heterogeneity and functional diversity of individual cells in a cell population [9–12].

We have previously developed and released mitoXplorer [13] and its first update, mitoXplorer 2.0 [14], a web-based platform to analyse and integrate -omics data in a mitochondria-centric way. At the heart of mitoXplorer are manually curated and annotated interactomes available for 4 species (human, mouse, fruit fly and budding yeast), which encompass all mitochondria-associated genes (mito-genes), as well as cytosolic genes important for mitochondrial function, such as glycolysis, Ca2+ signalling, nuclear transcriptional regulation of mito-genes, or apoptosis. These mito-interactomes are used to integrate bulk -omics data emanating from RNA-seq or proteomics studies. mitoXplorer 2.0 offers 9 interfaces for mining and integrating the data, allowing to analyse a single dataset, perform comparative analysis, time-series analysis, cross-species analysis, or population-based analyses. Moreover, with the 2^nd^ release, we made it possible to search for transcriptional, or signalling regulators of mito-genes. So far, mitoXplorer cannot be used for analysing single-cell -omics data. Single-cell based techniques are advancing at an unprecedented speed and will become more and more prominently applied in the future [10,15], opening new perspectives on understanding cellular states.

Here we introduce an update of the mitoXplorer web-tool, mitoXplorer 3.0, which we improved to work with sc/sn-RNA-seq data to uncover cell-type specific mitochondrial variability, as well as to investigate mitochondrial expression dynamics at a single-cell level. To this end, we developed a python script that takes the annotated single-cell data structure as input, and generates on the one hand pseudo-bulk data for subsequent mitochondria-centric data mining using the classical interfaces of mitoXplorer. Secondly, a mito-gene specific single-cell matrix is created that can further be used with 3 novel interfaces to mine the single-cell expression data from a mitochondrial perspective, allowing to identify different mitochondrial metabolic states of cell types. We demonstrate the predictive power of mitoXplorer 3.0 on a time-resolved, high-resolution single-nuclei RNA-sequencing dataset of Spinocerebellar Ataxia 1 (SCA1)[16]. By using pseudo-bulk- and single-cell analysis on human and mouse data in mitoXplorer, we identify some mito-genes that could play a role in SCA1 and that could potentially be used as marker genes for subpopulations of cell types, such as the Ca2+ signalling gene *Efhd1* or the protein stability proteins *HSPA1A* and *HSPA1B*. We furthermore uncover deregulated mito-genes in Purkinje cells in SCA1 mice that could contribute to the degeneration of this cell type, such as the glycolysis gene *AldoC*, which is downregulated already at an early stage of the disease, as well as the Ca^2+^ metabolism genes *Itpr1 and Slc25a13*.

## MATERIAL AND METHODS

### mitoXplorer 3.0 for analysing single-cell and single-nuclei transcriptome data

Mitoxplorer 3.0 is a further development of our mitoXplorer 2.0 pipeline allowing mitochondria-centric analysis of single-cell and single-nuclei sequencing data. We had two possible biological scenarios for analysis in mind (Figure 1). First, we wanted to take advantage of cell-type specific information originating from single-cell sequencing data. To this end, we wanted to generate cell-type specific pseudo-bulk data that can be analysed using standard differential expression analysis pipelines such as DESeq2 [17] or edgeR [18]. And second, we wanted to analyse mitochondrial variability within individual cell types and identify mito-gene based cellular subtypes. As single-cell data are typically large R-objects or h5ad files, we developed a pre-processing pipeline, scXplorer, which generates both the pseudo-bulk file of cell types, as well as the single-cell transcriptome data file specific for mito-genes. As input to scXplorer, the user only needs to provide the pre-analysed single-cell data object, containing the cell annotations and the UMAP coordinates.

**Figure 1:**
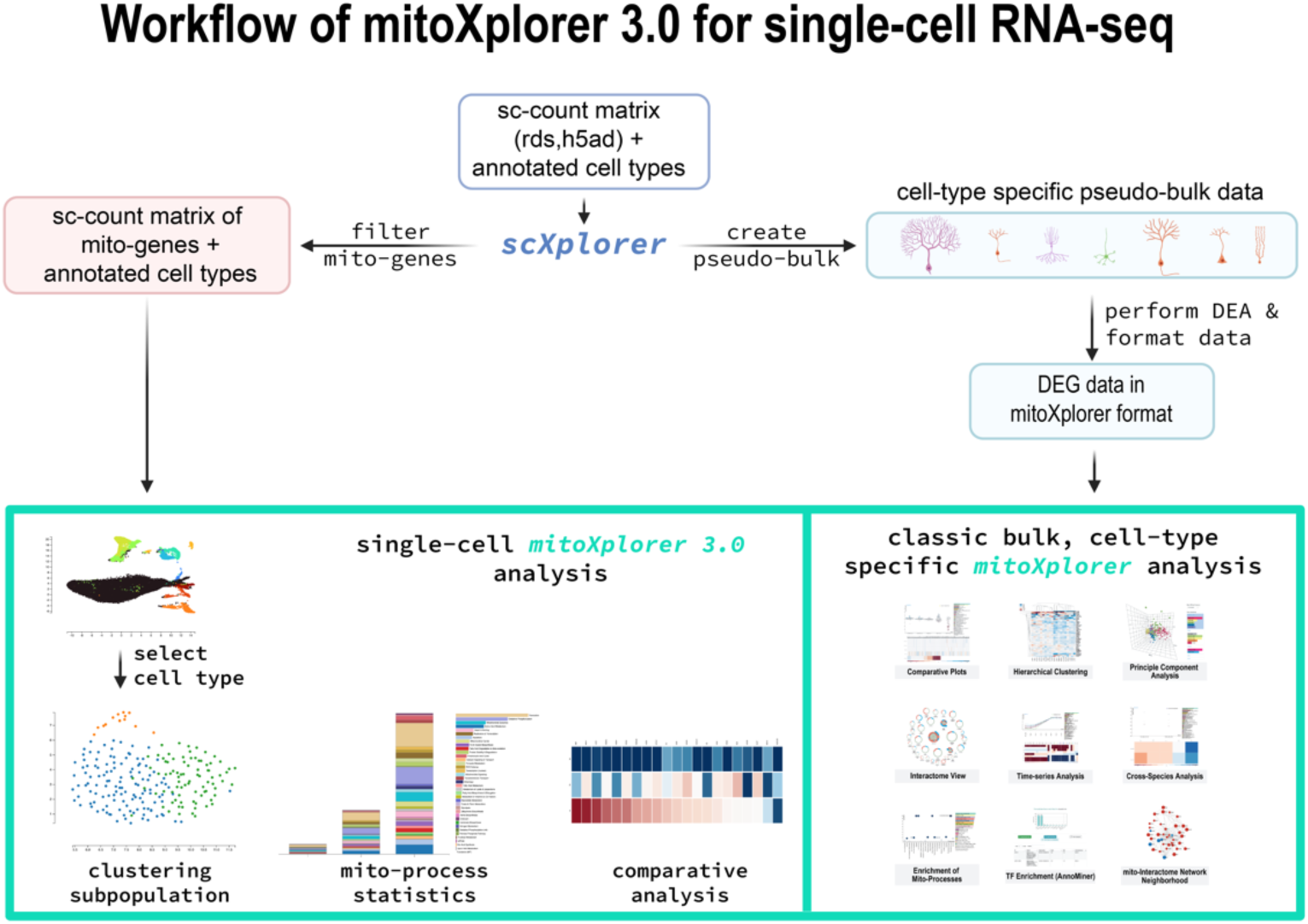
Workflow of single-cell RNA-seq data pre-processing and analysis with mitoXplorer 3.0. Single-cell RNA-seq data need to be pre-processed prior to mitoXplorer 3.0 analysis. The user needs to provide a scRNA-seq data object with annotated cell types in h5ad or rds format. For pre-processing the data, the scXplorer script was developed, which on the one hand creates pseudo-bulk data for annotated cell types, and performs differential expression analysis using pyDESeq2 [19]; and secondly, mito-genes as stored in the mitoXplorer database are selected from the scRNA-seq data structure for subsequent sub-clustering of cell types. Cell-type subpopulations can be analysed in the mitoXplorer platform for mito-processes present in the subpopulations, as well as on a mito-process basis for differential gene expression on the level of mito-processes.

### The scXplorer pre-processing script for scRNA-seq and snRNA-seq data objects

scXplorer was coded in Python 3.11. It requires an analysed and annotated single-cell sequencing data object as input, which has already undergone user-specified filtering, cell assignments, as well as generation of UMAP coordinates. The raw read counts are used for further processing with scXplorer. We recommend the usage of already existing software packages to analyse scRNA-seq/snRNA-seq data, such as Seurat [19] or scanpy [20], (for a typical workflow see [21]). When using the R-based Seurat package for filtering, cell assignments and UMAP generation, the resulting R-object should be saved as an RDS-file. Alternatively, the h5ad file format from the Anndata python [22] package can be directly used. Both can be fed into the scXplorer script. When the scXplorer pipeline is downloaded for local usage, the mito-gene set for filtering genes for single-cell based mitoXplorer 3.0 analysis is provided as part of the scXplorer package and needs to be localized in the folder of the executable script.

The command line to execute the formatting script is as followed :

~~~
python3 scXplorer.py -f [h5ad or rds file] -o [organism] -a [annotation] -n [output name]
~~~

parameters:

- `f [rds or h5ad file] :` specify the path to input rds or h5ad file(s)
- `o [organism] : specify the organism of the input file` Choose one of the following 4 organisms : human, mouse, fly and yeast.
- `a [annotation] :` specify the variable that stores the annotation of cells (mandatory for h5ad files).
- `c [control file]:` specify the control (wild-type) inputfile if there are two input files and you want to compare mutant and wild type.
- `g [genotype]:` alternatively, if mutant and wild type are in one file, specify the variable storing the genotype.
- `n [output name] :` specify output file name (optional).
- `XX [ cut-off ] :` specify cut-off of read counts for single-cell filtering (default: 30).

The script generates a folder named “*scXplorer”*, containing the mitoXplorer-ready, pre-processed and analysed pseudo-bulk data as well as the pseudo-bulk read counts deduced from the h5ad-formatted data file (in csv format). In addition, it generates the mito-gene filtered single-cell read count matrix in h5ad format, which can directly be uploaded to mitoXplorer (in zip format).

#### Filtering out genes with low read counts of and of mitochondria-encoded genes

To avoid the presence of genes with very low read counts for single-cell based comparative analysis, we chose to introduce a filtering step. The default value for raw read count filtering across all subpopulations is currently set to 30. We note here that mitochondria-encoded genes are not further filtered out of the dataset, though we ignore these genes in analysis of single-nuclei sequencing data.

#### Pseudo-bulk data pre-processing and analysis

The scXplorer script generates two csv files: The one with the extension “_pseudo_bulk.csv” contains pseudo-bulk data derived from the single-cell sequencing datafile: each annotated cell type is extracted, summarizing the read counts per gene over all cells of this cell type; at the same time, a pseudo-means is generated, which reflects the mean of the read counts of a gene over all contained cell types. Three pseudo-replicates are generated for each cell type and the pseudo-mean, to allow analysis as bulk RNA-seq data using all standard differential expression analysis packages. Pseudo-replicates are calculated by utilizing 33% of cells for each cell type, and the mean value for each gene is computed per replicate. The second csv file with the extension ‘_bulkXplorer.csv’ is the ready-to-use file for upload to mitoXplorer and for mining with the classical mitoXplorer 2.0 interfaces. Depending on the provided input, it will either contain a comparison of each cell type against the mean over all cell types (if only one single-cell RNA-seq dataset is provided), or contain data from the comparative analysis between two submitted RNA-seq datasets (for instance a wild-type and a mutant condition). To generate the mitoXplorer pre-analysed file, differential expression analysis is performed using the pyDESeq2 package [23], to obtain the log2 fold change (log2FC), and the p-value.

### Analysis of mito-gene based subpopulations in mitoXplorer 3.0

All analyses were done using Python v3.11 and scanpy [20]. All novel visualizations introduced to mitoXplorer 3.0 were developed using the D3.js library [24]. For single-cell analysis, the single-cell count matrix is filtered for mito-genes, which are exclusively used for all downstream analysis steps in the mitoXplorer 3.0 platform.

#### Clustering

Once the mito-gene filtered single-cell data is uploaded to the webtool, the user first needs to choose a cell type of interest, which will be clustered using the Leiden algorithm to identify subpopulations according to mito-gene expression profiles. The clustering resolution can be chosen by the user (default: 0.8). Clustering can be repeated at any time, choosing another clustering resolution or another cell type. Resulting clusters of subpopulations are displayed in a UMAP for visual inspection by the user. In case of two submitted conditions, the user can toggle the UMAP for displaying subpopulations, or conditions.

#### Differential expression analysis

To identify how much one subpopulation differs from the others, a differential expression analysis is performed using pyDESeq2 [23]. One subpopulation is compared to the mean of all other subpopulations excluding itself, calculating the log2FC and the p-value for each gene.

#### Mito-process based analysis and data visualization

Each subpopulation can be analysed according to mito-processes contained. Based on a chosen threshold of the p-value as determined by the differential expression analysis, genes that are below this threshold and are associated with mito-processes are counted for each subpopulation. This gives the user a global overview of the diversity of mito-processes present in each subpopulation.

### New visualization interfaces for mitoXplorer 3.0 single-cell RNA-seq data analysis

We have developed several novel interactive and user-friendly interfaces to mine single-cell RNA-seq data in mitoXplorer 3.0. To visualize subpopulation gene counts in mito-processes, we offer 3 interfaces: 2 stacked bar charts, and a heatmap coloured according to the number of genes in each mito-process for each subpopulation. Hovering over a box in this heatmap will display the number of genes found in the mito-process and subpopulation. Second, we offer to visualize each mito-process with an interactive, and sortable heatmap. Each gene for each subpopulation in the heatmap is coloured according to the log2FC. As is standard in all mitoXplorer interfaces, hovering over a gene box will reveal the information of the gene in the info-box at the left panel of the web-tool. The log2FC, the p-value, as well as the raw read counts are displayed in a pop-up box of the chosen cell population.

For human data, we additionally offer to analyse the distribution of subpopulations according to the cell cycle state, using the genes in S and G2M phases from Harmonizome [25] present in the mito-gene set. A prediction of the cell cycle phase for each subpopulation is given by distribution of the sum of scores for each cell cycle phase for the cells of the subpopulation.

Usage instructions of the new interfaces and on mining single cell data available in mitoXplorer 3.0 and used in this study can be found in Supplementary Material.

### MitoXplorer database

Pre-processed pseudo-bulk data were added to the mitoXplorer database and are freely accessible to the users. These currently include data from: the Tabula sapiens [26] and Tabula muris [27] projects, the Fly Cell Atlas [28] and datasets on spinocerebellar ataxia type 1 (SCA1) from [16] (GEO accession number GSE246183).

## RESULTS

### Use case: cell type- and cellular subpopulation-based mitochondrial variability of the SCA1 and WT data in human and mouse cerebellum

Neuronal diseases such as ataxias are especially profiting from single-cell or single-nuclei RNA-sequencing data, due to the cellular complexity and heterogeneity of the brain, as well as because of cell-type specific contributions to disease-onset, as is the case for most cerebellar ataxias (CAs). For introducing the novel features and predictive power of mitoXplorer 3.0, we therefore used data from Spinocerebellar Ataxia type 1 (SCA1). SCA1 is a genetic neurological disorder characterized by progressive loss of coordination and balance. It is caused by a polyQ-expansion in the *ATXN1* gene, leading to the production of the misfolded defective Ataxin-1 protein. Ataxin-1 is a transcriptional co-repressor and interacts with the fast transcriptional repressor, Capicua. The poly-Q expansion in Ataxin-1 leads to the loss of this interaction and the decrease of Capicua, resulting in the onset of SCA1 [29]. Ataxin-1 is also activating RORa dependent transcription [30], in conjunction with the transcriptional co-activator Yap/YapDeltaC [31]. Sp-1 has also been shown to interact with Ataxin-1 and there is experimental evidence that polyQ expansion of Ataxin-1 leads to deregulated Sp-1 target genes in Purkinje cells in mice [32]. Mitochondria play a crucial role in SCA1 and studies suggest that dysfunctional mitochondria may contribute to the degeneration of neurons in SCA1, especially in the Purkinje cells (PCs) and hippocampal neurons [33,34]: Mitochondrial impairment leads to increased oxidative stress and energy deficits, worsening the neurodegenerative process in SCA1. Understanding these mitochondrial implications could potentially offer new avenues for therapeutic interventions in the future.

We used pre-analysed and annotated mouse and human single-nuclei RNA-seq data of SCA1 from [16] (GEO accession number GSE246183). This dataset contains data from 10 human SCA1 patients, and 10 healthy control donors; as well as data from a mouse model of SCA1 knock-in (KI) containing the mutation *Atxn1*^154Q/+^, collected at different timepoints: 5, 12,18, 24 and 30 weeks of age. The phenotypes of these mice was described as follows [35]: motor coordination defects appeared as early as 5 weeks of age as assessed by poorer performance on the accelerating rotarod, together with a very subtle reduction of the total membrane area of the dendritic tree in Purkinje cells and growth retardation. From 12 weeks onwards, a reduced brain weight and reduced size of the fine dendritic arbour in PCs was reported, muscle wasting and an abnormal gate was noticed from 20 weeks of age onwards, together with impaired neuronal functions, curvature of the spine and atrophy of the lower limbs towards 30 weeks of age. Mice that were older than 30 weeks showed significant loss of PCs, reduced size of dendritic arborization and premature death.

The annotated file in h5ad format was used as input to scXplorer, and pre-processed using ‘sub4’ as annotation variable. We generated single-cell data and pseudo-bulk data of SCA1 and WT alone, as well as comparing SCA1 versus WT. For comparative analysis of pseudo-bulk data, we compared human SCA1 versus healthy donor, and equally for mice for each time-point (SCA1 vs WT control). We also performed pseudo-bulk data comparison within each dataset, comparing each cell type against the mean over all cell types.

### Part 1: Cell-type specific pseudo-bulk analysis

Mito-gene expression is generally very variable between cerebellar cell types (Supplementary Figure S1). Moreover, the most vulnerable cell type leading to the onset of CA is the Purkinje cell, with is, though very large, also sparse in the cerebellum. This justifies the usage of single-nuclei sequencing to investigate cerebellar neurodegenerative diseases, such as CA.

#### Enrichment of Glycolysis and Ca2+ signalling and transport in Purkinje cells from 18 week old SCA1 mice

We first set out to search for the cause of dysregulation of Purkinje cells in SCA1 animals. To this end, we first compared the single-cell data from wild-type and mutant animals throughout the time-series, using the newly developed UMAP visualization of the clustered PCs from the two conditions. The most striking difference in the signature of mito-processes between conditions was found in Purkinje cells (PCs) from 18 week old mice, where we saw a very clear separation of the wild-type and SCA1 population (Figure 2a). A slight separation was also visible at 12 and 24 weeks, but at this time-point, less PCs were sequenced leading to lower resolution of the data (Supplementary Figure S2.1). Based on these results, we focused on the 18 week timepoints for further analysis (results from Purkinje cells at 12 and 24 weeks are discussed in Supplementary Materials).

**Figure 2:**
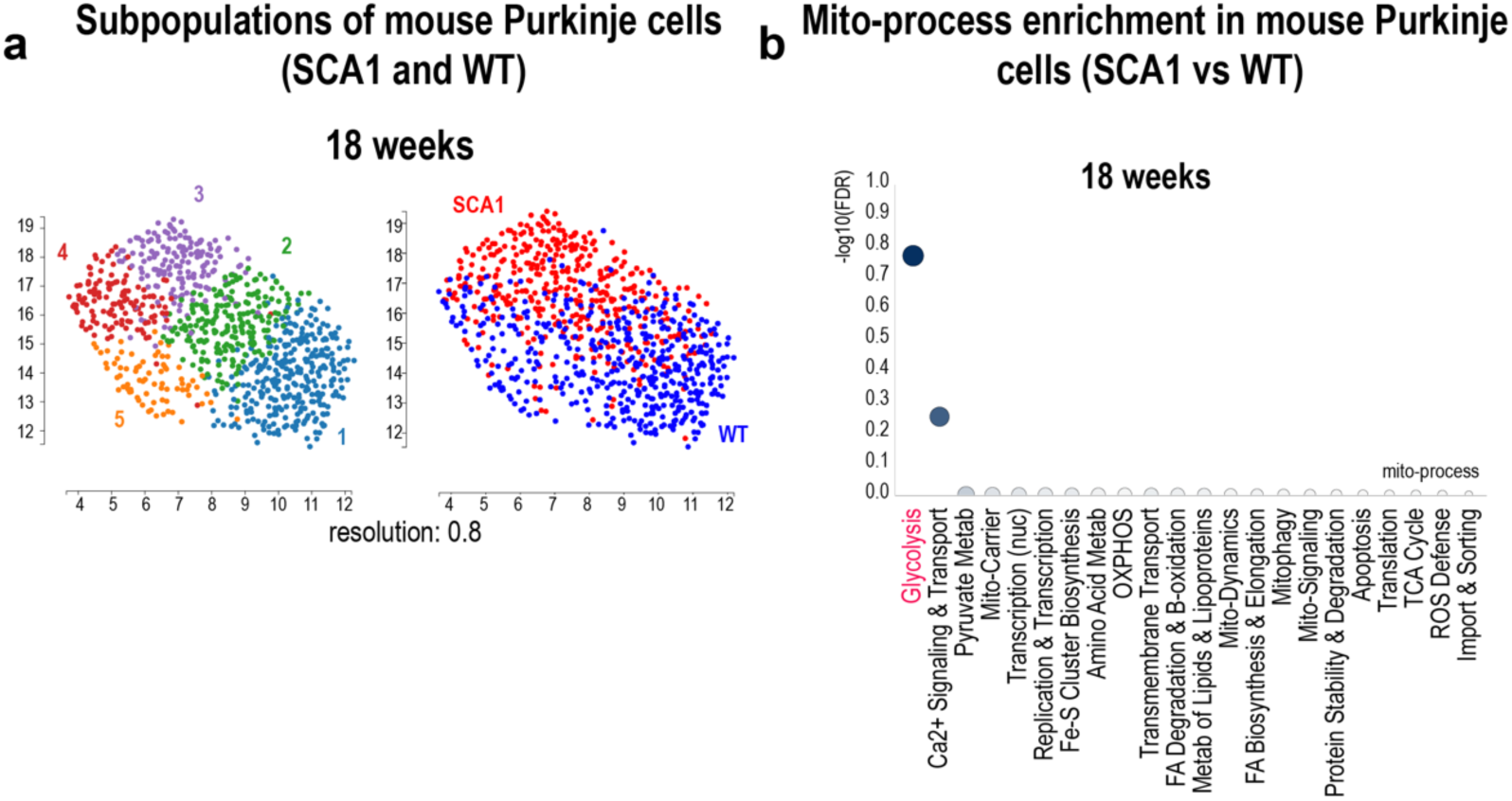
Glycolysis and Ca2+ signalling and transport are affected in SCA1 Purkinje cells. (a) UMAP of SCA1 and WT Purkinje cells and mito-centric overlay of single-nuclei populations from wild-type (blue) and SCA1 (red), isolated at 18 weeks. A clear separation can be seen between the 2 phenotypes. Cluster 3 (purple) is composed solely of SCA1 cells, while cluster 1 is predominantly WT cells, Cluster 2, 4 and 5 are mixed, with cluster 2 being slightly enriched in SCA1 and 5 in WT cells. (b) Enrichment analysis of 18 weeks PCs, comparing SCA1 to WT. Glycolysis and Ca2+ signalling and transport are the most enriched processes at 18 weeks.

Next we searched for enriched mito-processes in the pseudo-bulk data of 18 and 24 week PCs, comparing SCA1 to wild-type. At 18 weeks, we found that Glycolysis, as well as Ca2+ signalling and transport is significantly enriched (Figure 2b). Glycolysis was also enriched at 12 and 24 weeks (Supplementary Figure 3.1).

Based on these findings, we then searched for differentially expressed genes in these mito-processes in the pseudo-bulk of the time-series data. We therefore used the mitoXplorer time-series analysis and analysed the mito-processes Glycolysis (Figure 3a-c), as well as Ca2+ signalling and transport (Figure 3d-f).

**Figure 3:**
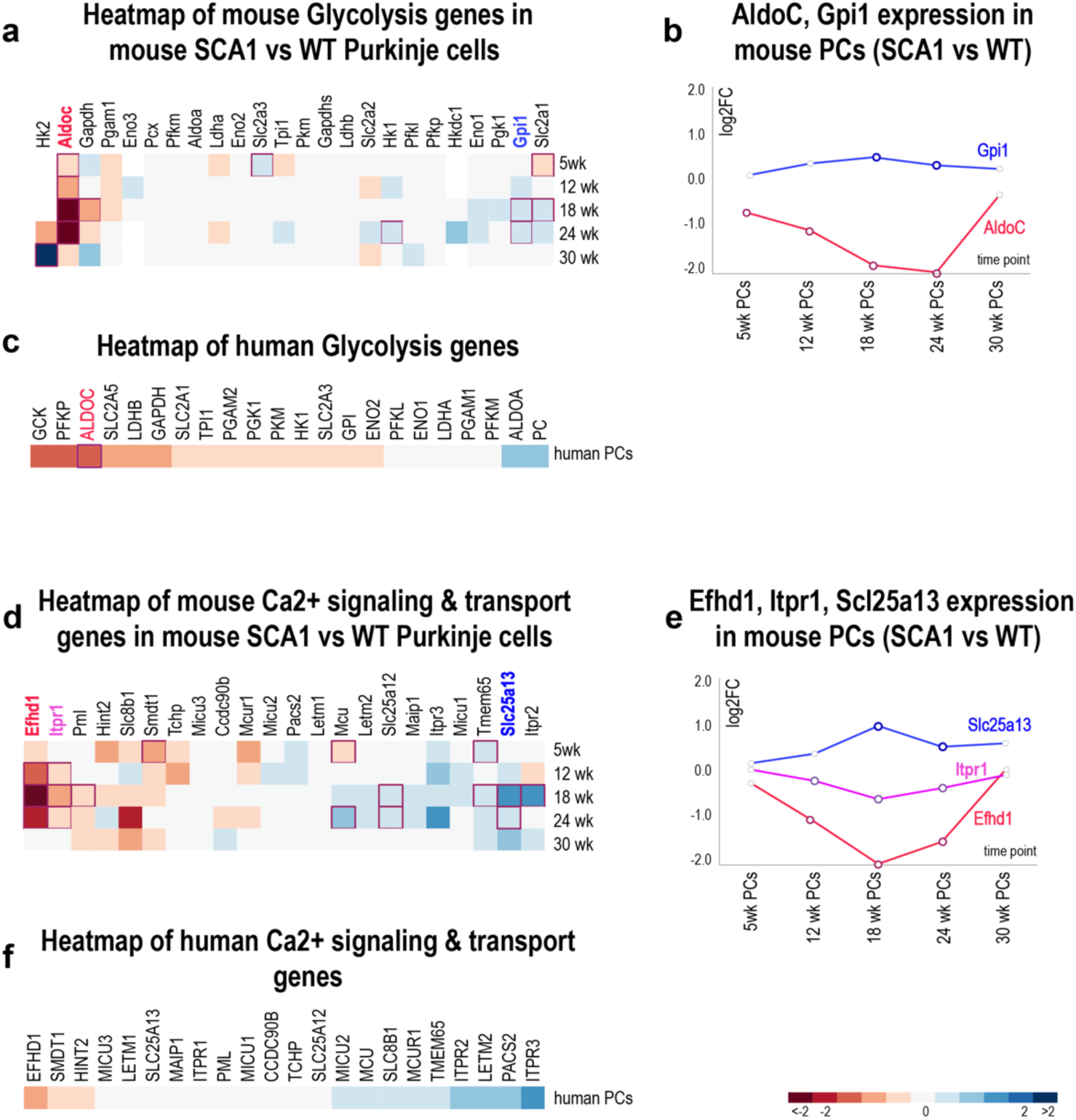
mitoXplorer 3 pseudo-bulk analysis comparing mouse Purkinje cells between SCA1 and WT. (a) Heatmap of Glycolysis of the mouse time-series comparison of SCA1 versus WT Purkinje cells. *AldoC*, as well as *Gpi1* are significantly altered in SCA1 versus wild-type. Highlighted genes are presented in 4b. (b) Time-series plot of *AldoC* (red) and *Gpi1* (blue) expression in SCA1 versus wild-type Purkinje cells. *AldoC* is strongly and significantly reduced at 12, 18 and 24 weeks. *Gpi1* is significantly induced at 18 and 24 weeks. (c) Heatmap of human Glycolysis genes of SCA1 patients versus healthy control. *ALDOC* is significantly reduced, while *GPI* is slightly, but non-significantly reduced. (d) Heatmap of Ca2+ signalling and transport of the mouse time-series comparison of SCA1 versus WT Purkinje cells. Several genes are significantly altered in SCA1 versus wild-type control at 18 and 24 weeks, including *Efhd1* (reduced), *Itpr1* (reduced), and the transporter *Slc25a13* (induced). (e) Time-series plots of *Efhd1* (red), *Itpr1* (pink) and *Slc25a13* (blue) expressions in mouse PCs comparing SCA1 to wild-type controls. (f) Heatmap of human Ca2+ signalling and transport genes in Purkinje cells of SCA1 patients versus healthy control. None of the genes are significantly deregulated, though *EFHD1* is downregulated with a p-value of 0.06. The colour bar indicates magnitude of down- (red) and upregulation (blue). Red lines around a box indicate a significant p-value (relevant for a,c,d,f), as do dark circles (in b,e).

In the mito-process Glycolysis, we found that *AldoC* is one of the genes that is significantly downregulated in PCs in SCA1, starting from 5 weeks of age (Figure 3a). This indicates that *AldoC* is affected early in the disease, even at a time-point when the phenotype is not yet manifested. The glycolytic enzyme Aldolase C converts fructose-1,6-bisphosphate to glyceraldehyde 3-phosphate (glyceraldehyde 3P, G3P) and dihydroxyacetone phosphate (DHAP)[36]. There are 3 isoforms, *AldoA*, *AldoB*, *AldoC*, whereby *AldoC* is specifically expressed in Purkinje cells, as well as other cell types of the brain [37].

The second gene that is significantly deregulated in SCA1 is the glycolytic enzyme glucose-6-phosphate isomerase (*Gpi1* in mouse and *GPI* in humans). *Gpi1* is a moonlighting enzyme. Within the cell, it is responsible for interconversion of glucose-6-phosphate and fructose-6-phosphate [38]. Extracellularly, its protein product is referred to as neuroleukin and it functions as a neurotrophic factor, promoting among others, the survival of sensory neurons [39].

The temporal dynamics of *AldoC* and *Gpi1* is shown in Figure 3b. *AldoC* was strongly downregulated at 18 weeks and remained low until 30 weeks *Gpi1* was on the other hand slightly induced at 18 and 24 weeks. It should be noted that at the 30 week time-point, when the phenotype of SCA1 was fully developed, very few PCs still remained, which could explain the lack of *AldoC* and *Gpi1* de-regulation at this time point [16]. *AldoC* (zebrin-II) has been used as a marker for PCs in several Ataxia studies [16,40]. In the data from human SCA1 patients, *ALDOC* was the only significantly deregulated gene in PCs in SCA1 patients in Glycolysis (Figure 3c). *GPI* was downregulated, however with a non-significant p-value.

The consistent and early downregulation of *AldoC* in PCs at all time-points makes it tempting to speculate that this gene is more directly affected by loss of Ataxin-1 dependent gene regulation, either as one early transcriptional target of the Ataxin-1 gene regulatory network, or as an indicator of an early metabolic shift upon loss of Ataxin-1 function. We therefore looked at *AldoC* expression in all cerebellar cell types, as well as at all time-points and found that it was consistently and - in most cases - significantly downregulated in cerebellar cell types at 5, 18 and 24 weeks in the SCA1 mouse, as well as human SCA1 patients (Supplementary Figure S2).

The second mito-process we investigated was Ca^2+^ signalling and transport. In this process, the mito-genes *Efhd1* and *Itpr1* were significantly downregulated at 12 and 24 weeks, while the transporter *Slc25a13* was significantly upregulated at 18 and 24 weeks (Figure 3d). Gene expression dynamics of these three genes are also shown in Figure 3e. *Efhd1* is a mitochondrial calcium ion sensor located at the inner mitochondrial membrane [41]. Itpr1 is an intracellular receptor for inositol 1,4,5- trisphosphate and upon stimulation, mediates Ca^2+^ release from the ER [42] and together with mtHsp70 and VDAC1 controls Ca^2+^ uptake of mitochondria [43]. *Itpr1* has been directly implicated in several types of Spinocerebellar Ataxias, such as SCA15, SCA16 and SCA29 [44] and, in the MalaCards database (https://www.malacards.org/), even with SCA1. *Slc25a13* is a Ca^2+^ regulated mitochondrial electrogenic aspartate/glutamate antiporter, allowing efflux of aspartate and entry of glutamate and proton within the mitochondria as part of the malate-aspartate shuttle [45]. In human SCA1 patients, we do observe reduction of *EFHD1*, however non-significantly, while *SLC25A13* is unchanged, as is *ITPR1* (Figure 3f).

In conclusion, in mouse cerebellum, Purkinje cells are the most affected cells upon mutation of Ataxin-1 at 18 weeks of age and several genes in the mito-processes Glycolysis and Ca2+ signalling and transport are significantly altered in the disease condition. We speculate that these events most likely contribute to the loss of these cells in SCA1. Some of these genes have already been associated with spinocerebellar ataxias, or are already used as marker genes for determining Purkinje cell viability.

### Part 2: Single-cell based analysis to uncover cellular subpopulations of cell types

With mitoXplorer 3.0, we also provide the possibility to perform mito-gene centric sub-clustering of scRNA-seq data and to identify subpopulations of cell types based solely on mito-genes. In order to demonstrate the functionality of these novel features, we aimed to identify mito-gene based subpopulations of Purkinje cells in the snRNA-seq data from mouse cerebellum. Given the high degree of variability between PCs originating from WT and SCA1 at 18 weeks, we decided to focus on this time point and used the combined SCA1 & WT dataset, to overlay the two different conditions and search for genes whose expression differs not only between subpopulations of PCs, but also between the two different conditions. Based on the clustering, as well as the overlay of conditions shown in Figure 2a, we consider subpopulations 1 and 5 as mostly wild-type, subpopulation 3 as SCA1 and subpopulations 2 and 4 as mixed.

We first analysed the mito-process based gene counts in the different subpopulations and found that the mito-processes Ca^2+^ signalling and transport contained the most differentially expressed genes between the subpopulations (Figure 4a).

**Figure 4:**
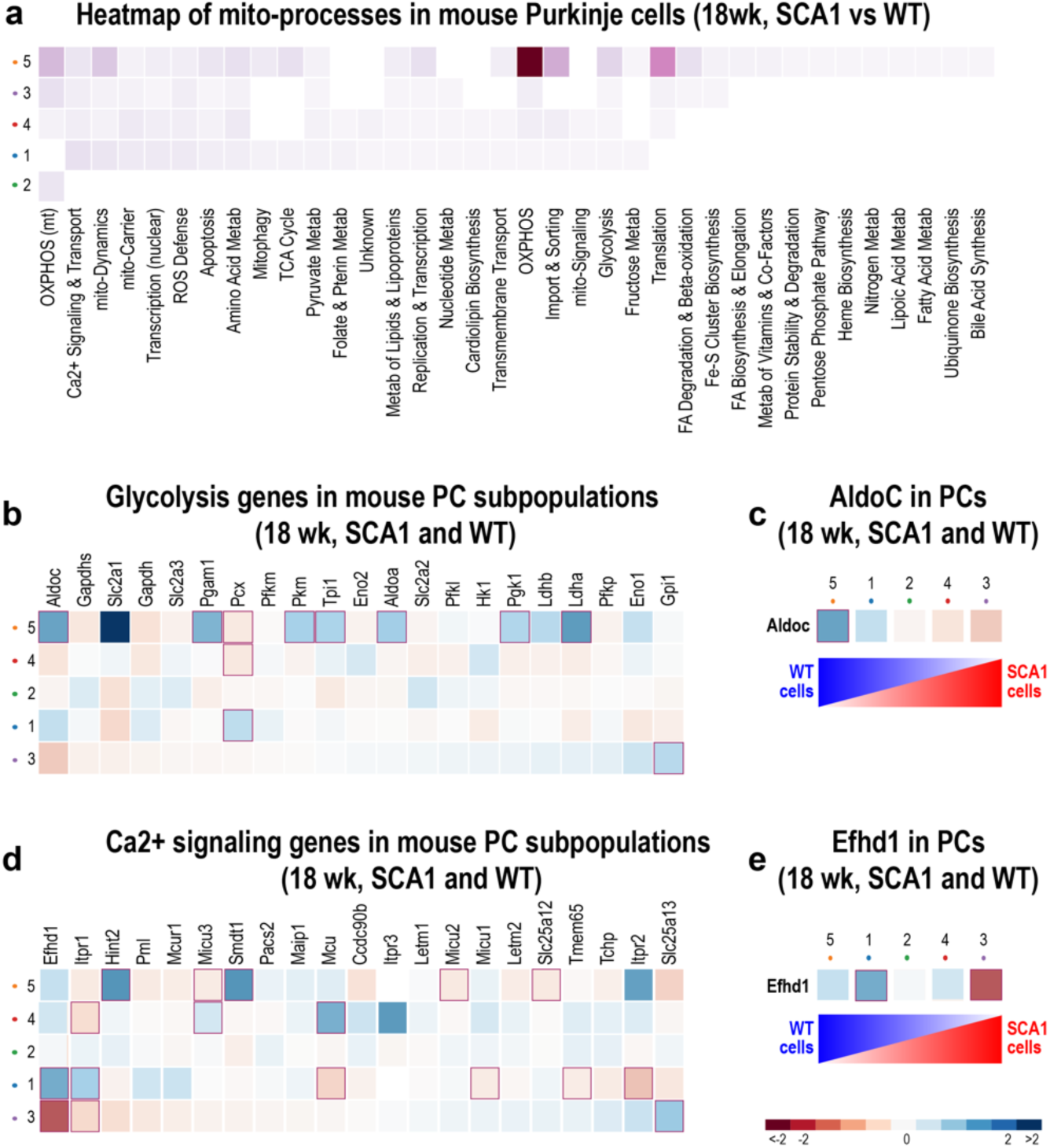
Single-cell mitoXplorer 3 analysis of Purkinje cells in 18 week SCA1 and WT cerebella. (a) Heatmap of mito-processes present in Purkinje cells in 18 week SCA1 and WT cerebella. Ca^2+^ signalling and transport, as well as Glycolysis genes are significantly different in the Purkinje cell subpopulations. (b) Glycolysis genes in mouse Purkinje cell subpopulations from SCA1 and WT. *AldoC* is highest and significantly upregulated in the subpopulations that consist mostly of wild-type cells (5, orange population in Figure 2a). Significant upregulation of *Gpi1* is observed in subpopulation 3 (lilac population in Figure 2a) that consists of SCA1 cells. (c) *AldoC* expression across the PC subpopulations. *AldoC* is upregulated significantly in subpopulation 5, mostly composed of WT cells and the lowest expression is found in subpopulation 3, composed of SCA1 cells. It is noteworthy that the p-value of downregulation in subpopulation 3 is 0.06 and thus, close to significance. (d) Ca^2+^ signalling and transport genes in mouse Purkinje cell subpopulations from SCA1 and WT. *Efhd1* is significantly downregulated in the SCA1 subpopulation (3, lilac) and upregulated in the WT subpopulation 1 (blue in Figure 2a), the gene *Itpr1* shows the same regulation, though to a slightly lesser extent. The transporter *Slc25a13* is significantly upregulated in the SCA1 subpopulation 3. (e) *Efhd1* expression in PC subpopulations. *Efhd1* is significantly downregulated in the SCA1 subpopulation 3, while it is high in subpopulations 1 and 5, which are mostly composed of WT cells.

Based on these data, as well as the enrichments of the pseudo-bulk analysis, we next decided to specifically look at the mito-processes Glycolysis and Ca^2+^ signalling and transport using the new mitoXplorer 3 single-cell analysis interfaces.

In the mito-process Glycolysis, we found *AldoC* to be significantly upregulated in subpopulation 5, containing mostly wild-type cells (Figure 4b, c). It is notable that there seemed to be a gradient of *AldoC* expression correlating with the relative numbers of WT Purkinje cells in the subpopulation (Figure 4c).

In the mito-process Ca^2+^ signalling and transport, the mito-gene *Efhd1* was significantly and strongly reduced in the SCA1 subpopulation of Purkinje cells, while *Itpr1* was mildly but significantly reduced, and *Slc25a13* was significantly upregulated in the SCA1 subpopulation 3 (Figure 4d). We further plotted the regulation of gene expression of *Efhd1* according to the composition of subpopulations and found again that there was a decrease in expression from the wild-type clusters 1 and 5, towards the mixed and SCA1 clusters 2, 4 and 3, respectively (Figure 4e).

In conclusion, using the newly developed interfaces of mitoXplorer 3.0 for single-cell analysis, we have identified significantly deregulated genes in the mito-processes Glycolysis and Ca^2+^ signalling and transport. This analysis confirmed our interest in the *AldoC* gene, as well as in *Efhd1*, which are downregulated in SCA1 cell populations and shed new light also on glycolysis and Ca^2+^ metabolic genes as potential markers for Purkinje cell loss in SCA1.

We finally investigated the Purkinje cell subpopulations in WT and SCA1 condition alone at 18 weeks of age (Figure 5). We detected 3 mito-gene centric subpopulations of Purkinje cells in WT (Figure 5a), as well as in SCA1 (Figure 5b), respectively. We looked at mito-processes with significant differentially expressed genes for the respective 3 subpopulations in the two conditions and could see that the profile of associated mito-processes has changed in SCA1 at this age. While in the wild-type population, most marker genes originated from oxidative phosphorylation (OXPHOS), in the SCA1 condition, ROS defence was the most prominent mito-process. Given these results, we were most interested in investigating the bioenergetic processes OXPHOS and Glycolysis in the two conditions. Subpopulation 3 of wild-type PCs showed an induction of OXPHOS genes compared to the other two populations, indicating that this is the metabolically most active subpopulation (Figure 5c). Interestingly, this is also the subpopulation with highest levels of *AldoC* (Figure 5e). For SCA1 cells, we found two glycerol-3-phosphate dehydrogenases in the mito-process OXPHOS that together form the Glycerol phosphate shuttle (GPS) significantly induced in subpopulation 3, namely *Gpd1* and *Gpd2* (Figure 5d). Consequences of the GPS are a reoxidation of cytoplasmic NADH to NAD+, a potential bypass of complex I due to electron transfer by *Gpd2* to CoQ, with concurrent reduction of FAD to FADH2 and finally, the availability of cytosolic metabolite glycerol-3-phosphate (G3P) that connects glycolysis, lipogenesis and OXPHOS. Furthermore, the mitochondrial GPD2 gene could be involved in elevated production of ROS (for a review, see [46]). It was demonstrated that protein levels of respiratory chain (RC) components were reduced in these SCA1 mice, together with RC activity in disease vulnerable Purkinje cells and elevated oxidative stress [33]. Upregulation of the GPS could therefore be a reaction to loss of Complex I function to restore RC function in the cells. We looked next on Glycolysis and found that in wild-type in subpopulation 3, *AldoC* is upregulated compared to the other 2 subpopulations, rendering it the *AldoC* positive PCs (Figure 5e). Next to *AldoC*, several other glycolytic enzymes are upregulated, suggesting that this is the metabolically most active PC subpopulation. Finally, in SCA1, only the transporter *Slc2a1* was significantly differentially expressed in Glycolysis. We detected very few reads of *AldoC* and the difference in expression between the 3 subpopulations was not significant (Figure 5f).

**Figure 5:**
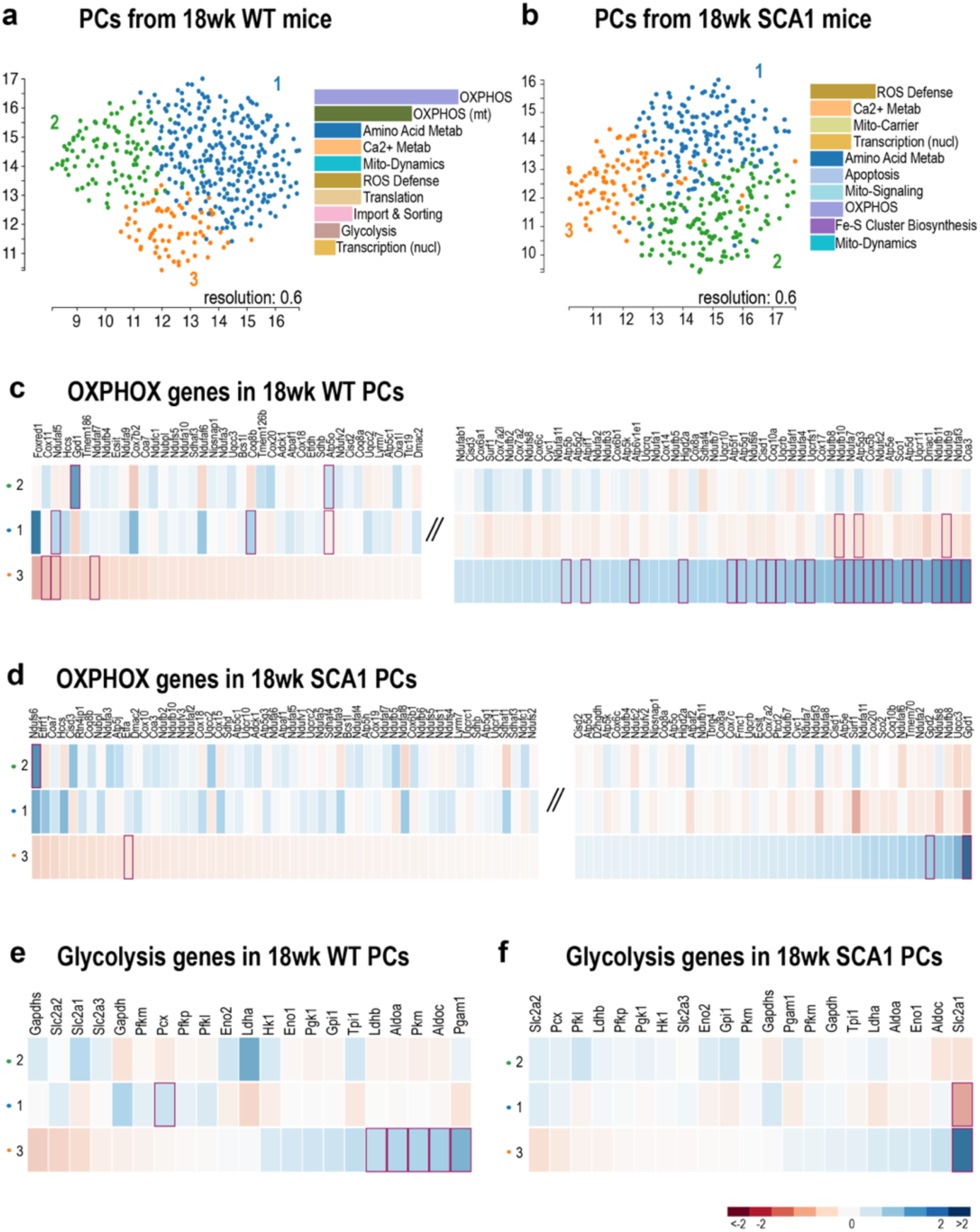
Single-cell mitoXplorer 3 analysis of Purkinje cells in 18 week SCA1 mice. (a) UMAP and mito-processes most enriched in wild-type, 18 week Purkinje cells. At a resolution, we found 3 subpopulations. Genes from the mito-processes OXPHOS, Amino acid metabolism, Ca2+ metabolism, but also Glycolysis were most differentially expressed. (b) UMAP and mito-processes associated in 18 week SCA1 Purkinje cells. Again, 3 populations were found at a resolution of 0.6. The mito-processes associated with the three cell populations have changed, with ROS defence having the most differentially expressed genes, followed by Ca2+ metabolism and mitochondrial carrier.(c) OXPHOS genes in 18 week wild-type PCs. OXPHOS harboured most of the significantly differentially expressed genes in WT PCs, with many genes upregulated in subpopulation 3, which is also the *AldoC* positive subpopulation. (d) In SCA1 PCs, in accordance with a potential elevated oxidative stress, the genes *Gpd1* and *Gpd2*, which form the glycerol phosphate shuttle that links OXPHOS, Glycolysis and lipid metabolism are significantly elevated in subpopulation 3. (e) Glycolysis genes in 18 week wild-type PCs. The genes *Pgam1*, *AldoC*, *Pkm*, *AldoA* and *Ldhb* were induced in subpopulation 3, which can be described as the *AldoC* positive cell population. (f) Glycolysis genes in 18 week SCA1 PCs. Only the glucose transporter *Slc2a1* was significantly deregulated, with highest expression subpopulation 3. *AldoC* is no longer significantly differentially expressed in SCA1 PCs in any of the subpopulations.

Taken together, by analysing the subpopulations of WT and SCA1 Purkinje cells separately, we identified a metabolically active subpopulation 3 in wild-type, which seemed to have shifted gene expression patterns in SCA1, expressing lower amounts of OXPHOS genes and losing also *AldoC* expression in the metabolically more active subpopulation.

Following the example of our analysis of 18 week mouse Purkinje cells presented above, we performed a comprehensive analysis of all mouse and human cell types of this SCA1 dataset, looking for those cells that show a marked difference between wild-type and mutant in single-cell sub-clustering. The results of this analysis are discussed in the section *Part 3* in Supplementary Material, and are shown in Supplementary Figures S3 and S4.

## DISCUSSION

We here introduce an update of the mitoXplorer 2.0 web-tool that has been adapted to analyse single-cell RNA-seq data from a mitochondria-centric perspective. We have developed and make available a pre-processing script, scXplorer that pre-processes pre-analysed and cell-type annotated scRNA-seq/snRNA-seq data and generates ready-to-be used data for upload and mining with mitoXplorer 3.0.

ScXplorer generates on the one hand cell-type specific pseudo-bulk data from an annotated single-cell or single-nuclei sequencing file. Using scRNA-seq/snRNA-seq data for a cell-type specific pseudo-bulk analysis carries immense power as it can reveal a cell-type expression dynamics [47,48]. This is especially true in highly complex and heterogeneous tissues, such as the brain. However, also other tissues are rarely composed of a single cell type. Focusing on a mito-centric analysis using the mitoXplorer 3.0 web-tool enables users to also mine cell-type specific pseudo-bulk data easily with respect to mitochondrial features. Cell-type specific mitoXplorer 3.0 analysis will help understand mitochondrial expression dynamics on a cell-type level and to identify cell-type specific metabolic differences in heterogeneous tissues.

On the other hand, our script also generates a mito-gene single-cell count matrix which can be used to perform true single-cell based analysis from a purely mitochondrial perspective. This allows to sub-cluster cell types based on mitochondrial features. With novel interactive data mining interfaces, mitoXplorer 3.0 further allows to analyse subpopulations based on mito-processes and identify differentially expressed genes in the different populations.

We have used data from spinocerebellar ataxia type 1 (SCA1) to demonstrate the new features of mitoXplorer 3.0 and to assess the mitochondrial component to the loss of Purkinje cells in SCA1. We explored the mito-gene expression landscape of Purkinje cells in this disease. The published data were furthermore giving us the possibility to explore the temporal mitochondrial expression dynamics of SCA1 development in mouse cerebellum. Using the pseudo-bulked data and the classical interfaces, we could find that *AldoC* is one of the genes that is majorly affected in PCs throughout the progression of the disease. Not only is it downregulated already at an early stage of the disease, when the phenotype is not yet manifested. But this gene is also downregulated in most other cell types of the diseased cerebellum we analysed in mouse, as well as in human, making it an early response gene to the loss of Ataxin-1. It was recently published that SCA1, SCA2 and SCA7 all show loss of *AldoC* (zebrin-II) in Purkinje cells at the onset of the disease [40]. Ataxin-1, Ataxin-2 and Ataxin-7 are all transcription factors and despite different biological functions, the signature of CAG- polyQ mutated versions of these three proteins result in overlapping gene expression signatures in the cerebellum [49]. While a direct regulation of *AldoC* by Ataxin-1 cannot be excluded, the shared loss of expression of this gene could indicate an early sign of a metabolic shift that is common to these types of cerebellar ataxias. Together with the decrease in *AldoC* expression in PC subpopulations, we also observed less upregulation of OXPHOS-related genes in this subpopulation, indicating that there is a general decline in metabolic activity in this PC subtype.

Another gene we identified using pseudo-bulk analysis of the dataset is the Ca^2+^ signalling and transport gene *Efhd1*. *Efhd1* is supposedly involved in regulation of so-called mito-flashes in a Ca2+ dependent way [41]. Mitoflashes are stochastic mitochondrial events that include bursts of superoxide production, transient depolarization of the mitochondrial membrane potential and reversible opening of the membrane permeability transition pore [50]. These mitoflashes are highly conserved and therefore represent most likely part of the basic functional repertoire of these organelles. *Efhd1* knockout mice show reduced levels of ROS, reduced mitoflash events in cardiomyocytes [51], as well as resistance to hypoxic injury. In dorsal root ganglia, the knockout of *Efhd1* led to reduced axonal growth *in vitro* and enhanced cell death together with mitochondrial dysfunction and a decrease in axonal ATP levels [52]. The role of *Efhd1* in Purkinje cells remains to be elucidated. Its downregulation, whether through Ca2+ regulated mitoflashes, or other mitochondrial stress signals, could lead to PC dysfunction and decline. It would be interesting to test whether mitochondrial mitoflashes are part of the pathophysiology, or the response to it, in PCs in SCA1. While the original work from Tejwani and colleagues provided a comprehensive and thorough analysis of the single-nuclei data from SCA1 animals and human patients on genome-scale level, this more focused and in-depth analysis of mitochondria-associated processes and genes using mitoXplorer 3.0 provides a complementary analysis, focusing on the role of mitochondria in this disease.

There are some inherent problems when using scRNA-seq/snRNA-seq data that should be further considered. First, these types of data will often result in low read counts for a number of genes per cell, bordering on noise due to sequencing artifacts. As mitoXplorer is focusing on a specific set of genes, this problem is enhanced. We observed this problem not only for single-cell based analysis, but in some cases also for pseudo-bulk analysis. For this reason, we filter for low expressed mito-genes by using cut-offs for both, differential expression analysis of single-cell as well as pseudo-bulk data. Furthermore, the read counts, together with the log2FC and the p-value, should be considered by the user when interpreting the results. Additionally, we have implemented visualizing significant p-values which are below 0.05 in the heatmap display of both, bulk and single-cell interfaces in mitoXplorer 3.0. Second, when performing mito-gene centric sub-clustering of single-cell data in mitoXplorer, the user needs to carefully analyse the resulting UMAP and try different clustering resolutions. This is a critical step in single-cell data analysis and only when cell populations are well separated, further analyses will provide meaningful information. Finally, the user needs to provide a pre-analysed and annotated scRNA-seq/snRNA-seq dataset for subsequent pre-processing and mitoXplorer 3.0 analysis. This dataset should contain the annotated cell types. This process is highly subjective and the user will strongly influence the assignment of the cell type in single-cell experiments. Some tools exist that can help in cell type assignment [53]. We recommend performing this process very carefully to avoid later artifacts in analysing data with tools like mitoXplorer.

## Availability

mitoXplorer 3.0 is available as a web-server at: https://mitoxplorer3.ibdm.univ-amu.fr; and for download from our gitlab repository at: https://gitlab.com/habermann_lab/mitox3.

## Author Contributions:CRediT

MH: conceptualisation, data curation, formal analysis, methodology, software, validation, writing – original draft; AdB: formal analysis, validation, writing – reviewing and editing; HP: conceptualisation, validation, funding acquisition, writing – review and editing; BHH: conceptualisation, formal analysis, funding acquisition, methodology, project administration, software, supervision, validation, visualisation, writing – original draft, writing – review and editing.

## Acknowledgements

We thank Arnaud Mourier, Baptiste Libe-Philippot for critical reading of the manuscript. We thank the Habermann team for helpful discussion.

