## Supplementary Figures S1-S6 for "mitoXplorer 3.0, a web tool for exploring mitochondrial dynamics in single-cell RNA-seq data"

#### Supplementary Figure S1:

##### Heatmap of differential gene expression in cerebellar cell types of 24 week old mice

**FA degradation & beta-oxidation**

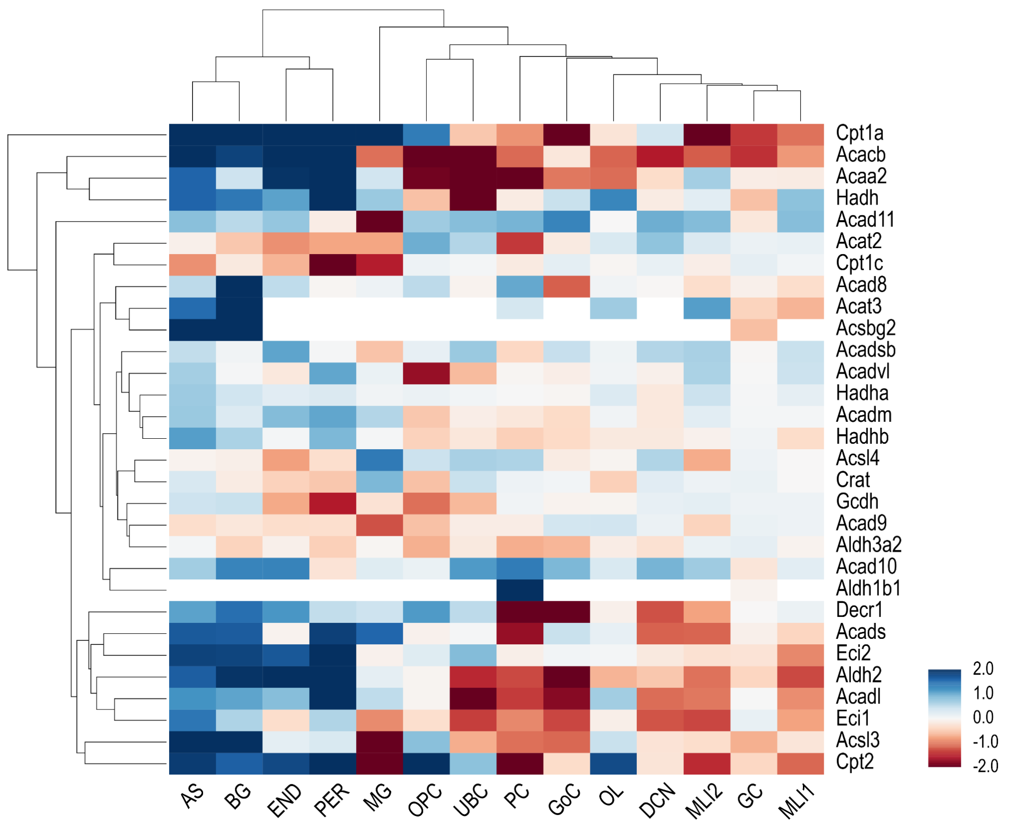


**Supplementary Figure S1: mitoXplorer 3.0 heatmap uncovers gene expression differences in cerebellar cell types.** We first wanted to investigate the cell-type specific heterogeneity of gene expression in this highly heterogeneous tissue. We therefore chose wild-type mouse data from 24 weeks old mice and used the heatmap function to visualize the differences in gene expression of each cell type against the mean over all cell types, using the pseudo-bulk data generated from the single-nuclei transcriptomic data. As shown in Figure 2 for the process *Fatty acid degradation & beta-oxidation,* there was a marked difference in gene expression of the different cell types in the cerebellum. This type of heterogeneity was seen for all mito-processes and thus, it is justified to use snRNA-seq data and perform differential expression analysis on pseudo-bulks of cell types to investigate highly complex and heterogeneous tissues, such as the cerebellum. As an example, we used the process *Fatty acid degradation & beta-oxidation* from 24 week old wild-type mice. Pseudo-bulks for each cell type were generated from single-cell data. Comparison was done of each cell type versus the mean over all cell types. GC - Granule cells; DCN - Deep cerebellar nuclei; UBC - Unipolar brush cells; PC - Purkinje cells; MLI1 - Molecular layer interneuron 1; MLI2 - Molecular layer interneuron 2; GoC - Golgi cells; AS - Astrocytes; BC - Bergmann glia; OPC - Oligodendrocyte progenitor cells; OL - Oligodendrocytes; MG - Microglia; PER - Pericytes; END - Endothelial cells

#### Supplementary Figure S2:

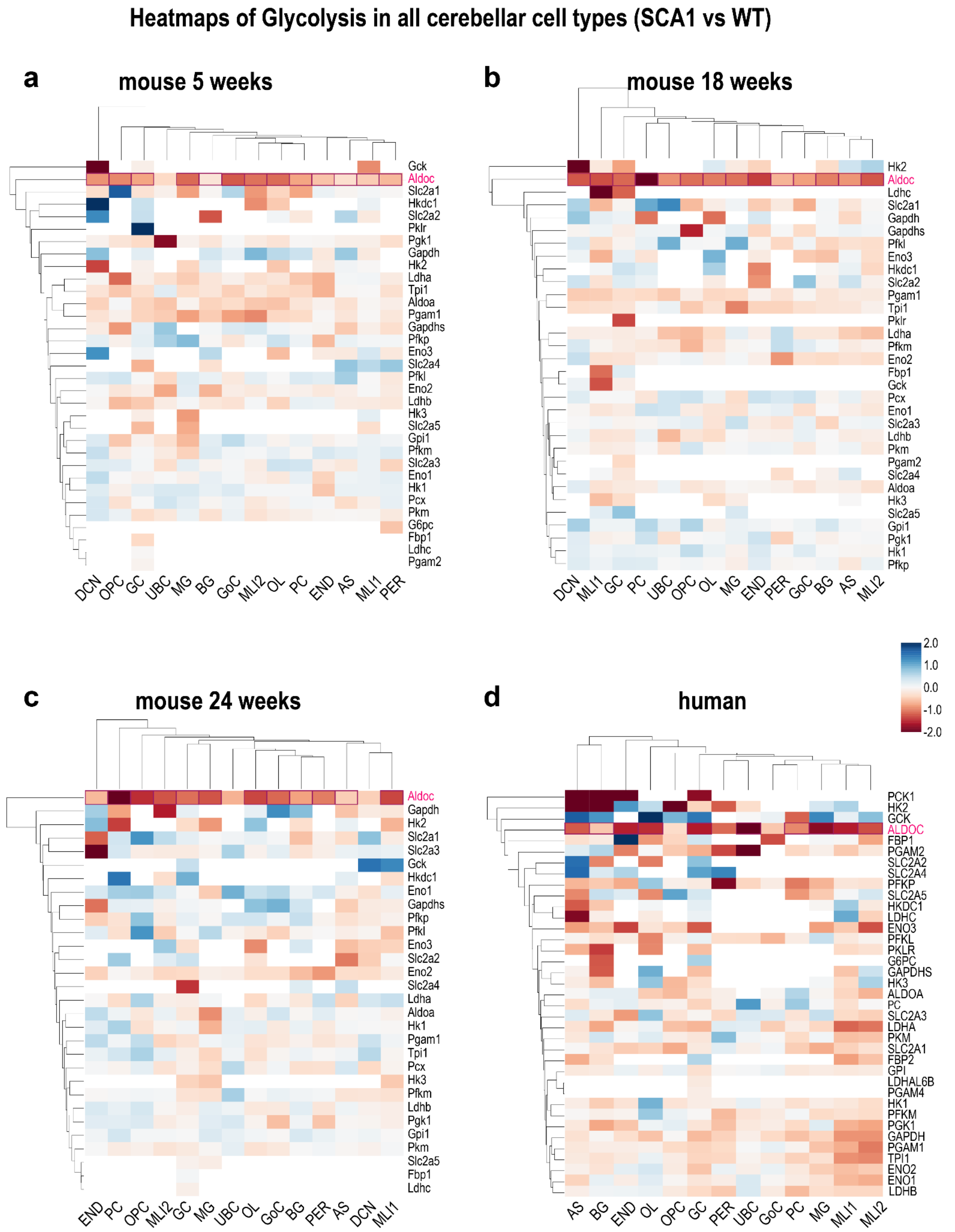


**Supplementary Figure S2:** ***AldoC* is reduced in all cell types at several time-points in SCA1 Purkinje cells.** Heatmap analysis of mouse 5 week (a), 18 week (b) and 24 week (c) data from cerebellum. (d) Heatmap analysis of all cell types in the human cerebellum. *AldoC/ALDOC* is significantly downregulated with some exceptions in all cell types in the datasets. Red lines around boxes indicate significant p-values. GC - Granule cells; DCN - Deep cerebellar nuclei; UBC - Unipolar brush cells; PC - Purkinje cells; MLI1 - Molecular layer interneuron 1; MLI2 - Molecular layer interneuron 2; GoC - Golgi cells; AS - Astrocytes; BC - Bergmann glia; OPC - Oligodendrocyte progenitor cells; OL - Oligodendrocytes; MG - Microglia; PER - Pericytes; END - Endothelial cells.

#### Part 3: Comprehensive analysis of mouse and human cerebellar cell types using mitoXplorer 3.0

In mice, we found 3 cell types to be most affected in SCA1 that showed separation on the UMAP between SCA1 and WT. Among those were Purkinje cells at 12 and 24 weeks (Supplementary Figure S1.1), which confirmed on one hand the downregulation of *AldoC* at 12 weeks in a SCA1 rich subpopulation; as well as the *Itpr1* gene at 24 weeks that was upregulated in WT-enriched subpopulations compared to others. We further found Deep cerebellar nuclei (DCN) with distinguishing UMAP distributions between the two conditions at 5 and 24 weeks (Supplementary Figure S1.2), which displayed interesting expression dynamics of Ca2+ signaling and transport genes at both time-points with downregulation of *Slc25a13* and *Itpr2* in SCA1 enriched subpopulations at both timepoints. Finally, Bergmann Gia (BG) at 18 weeks showed a differential distribution of SCA1 and WT cell on the UMAP that was however harder to separate. Nonetheless, we defined 2 subpopulations with slightly increased levels of SCA1 BGs. These cells had many mito-related processes enriched, including Glycolysis, Ca2+ metabolism and Amino acid metabolism and some of the same genes were differentially regulated as we found in other cell types, such as *Itpr1* which was higher in the predominantly WT population (Supplementary Figure S1.3 a-d).

The human data was much richer in cell numbers and contained in many cell types subpopulations that were specific to one of the two conditions (see Supplementary Figures S2.1-2.4). Cell types that showed separation on the UMAP included Astrocytes (AS), Bergmann glia (BG), Molecular layer interneurons 1 and 2 (MLI1 and MLI2), Oligodendrocyte precursors (OPC) and Oligodendrocytes. Purkinje cells could not be separated on the UMAP, which most likely reflects the very late state of the disease in the donor patients that is accompanied by loss of this cell type. One of the most striking observations we made was connected to the mito-process Protein stability and degradation, which was consistently and strongly enriched in nearly all cell types on pseudo-bulk and single-cell level analysis. When analyzing this process more closely, we identified the two Heat shock proteins Hsp70 1A and 1B, encoded by the *HSPA1A* and *HSPA1B* genes as significantly and strongly upregulated in SCA1, at least on RNA-level. We therefore further analyzed this process using the human pseudo-bulk data and could verify that these two genes were consistently, and in many cases significantly upregulated across all cell types (Supplementary Figure S3), probably indicating significant neurodegenerative stress in the cerebellum at this stage of the disease.

**Supplementary Figure S3.1:**

#### Analysis of subpopulation differences between selected mouse cerebellar cell types

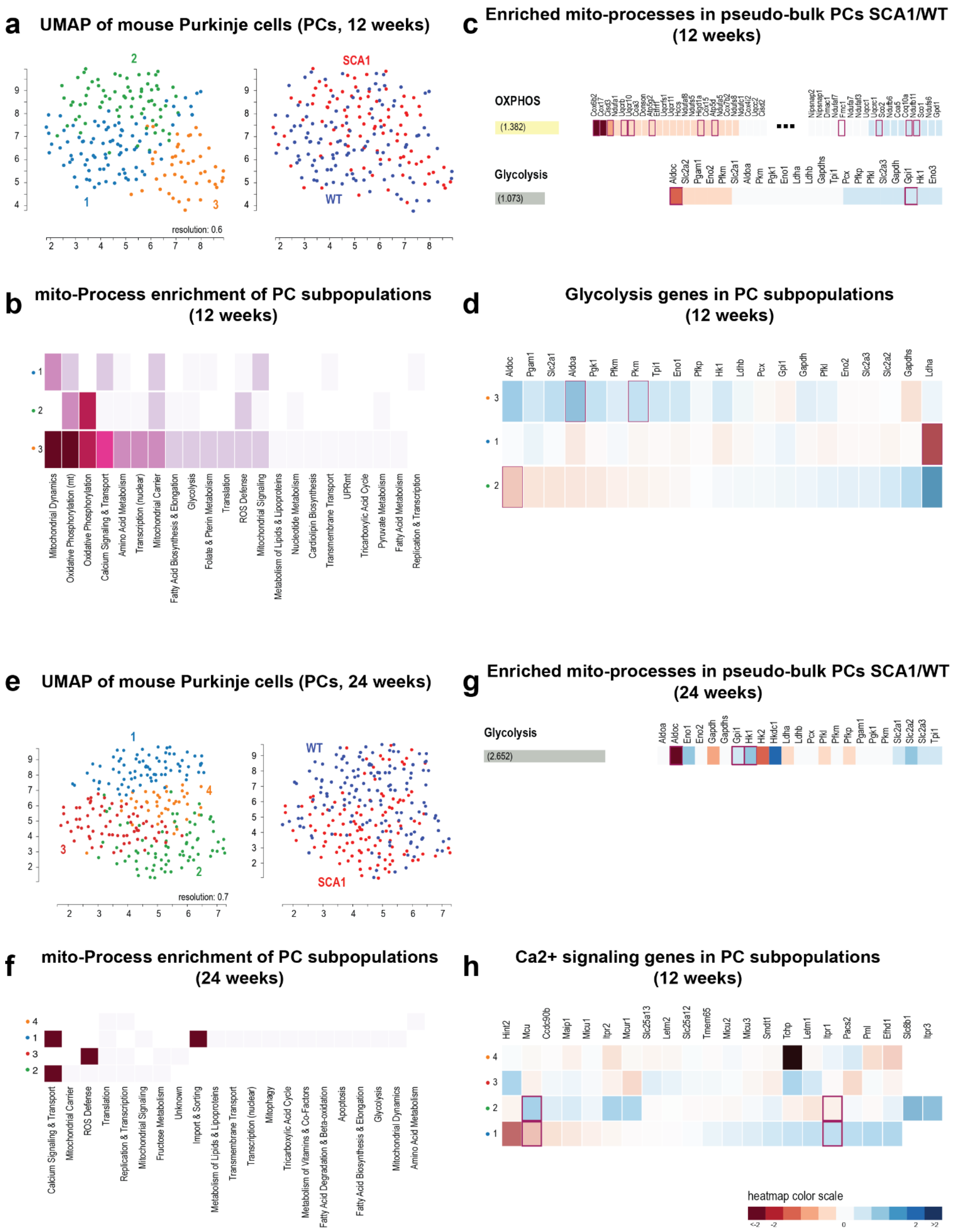

#### Supplementary Figure S3.2:

##### Analysis of subpopulation differences between selected mouse cerebellar cell types

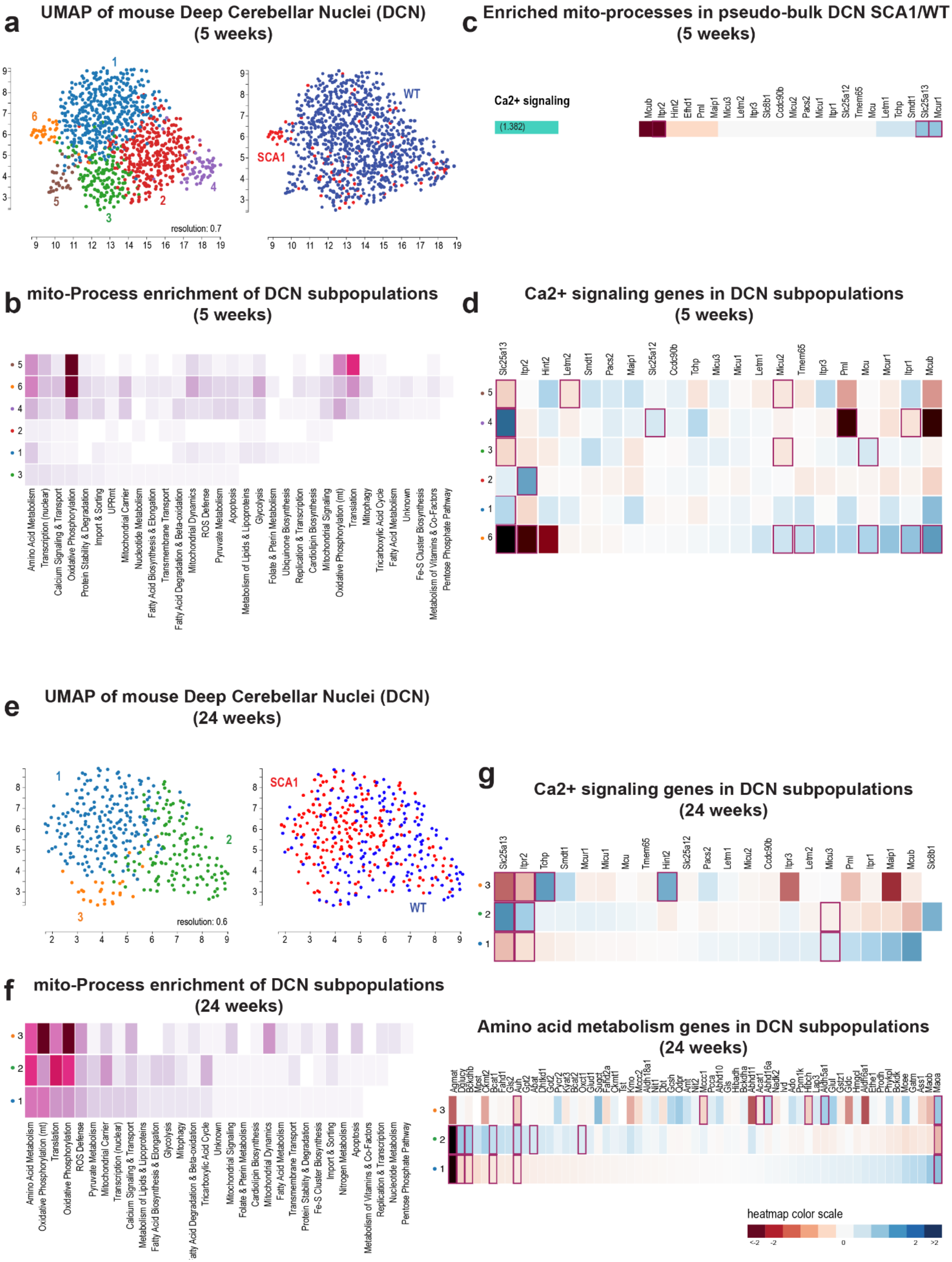

**Figure S3.3**

#### Analysis of subpopulation differences between selected mouse cerebellar cell types

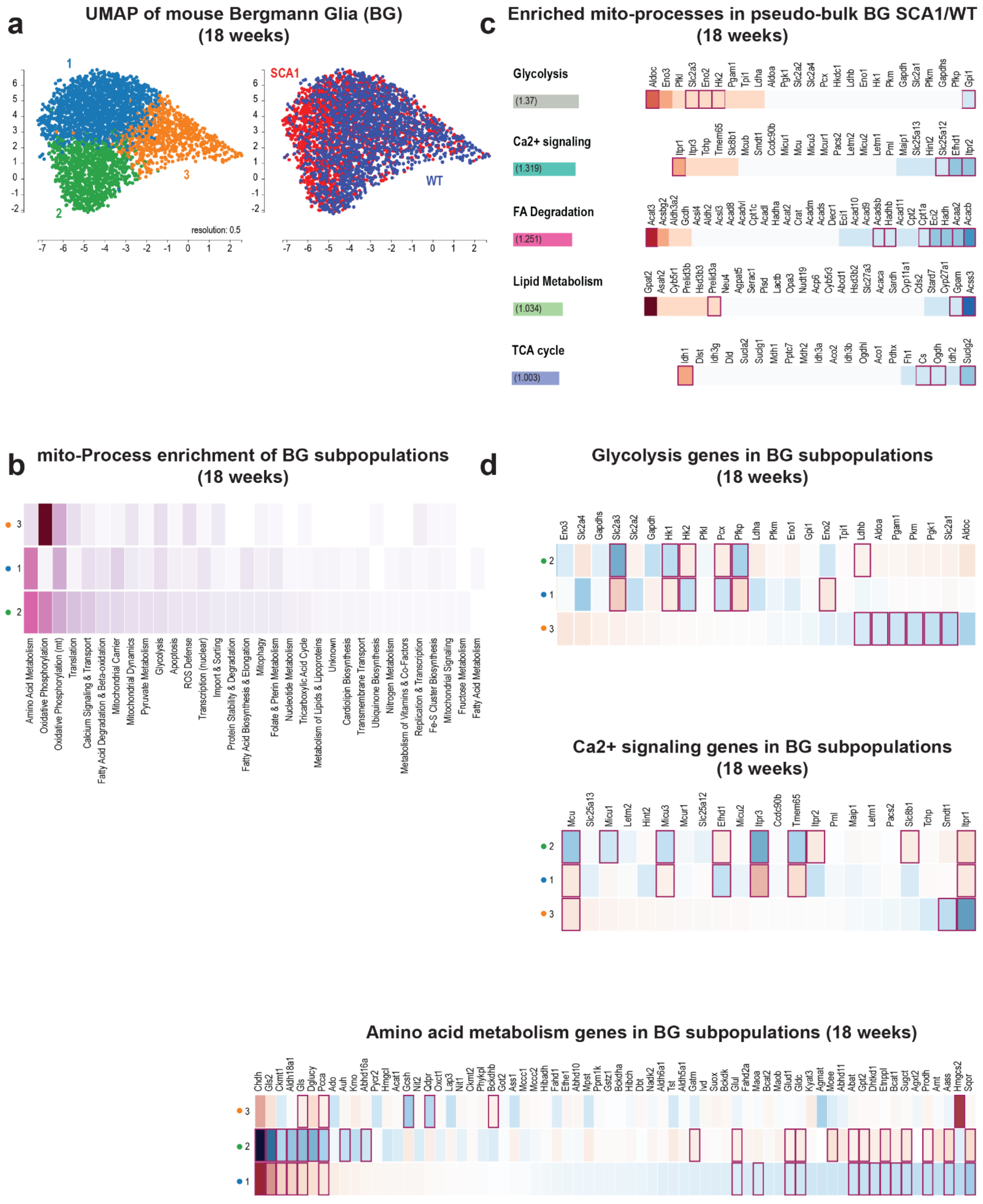

**Supplementary Figure S3: Single-cell and bulk analysis of mouse cerebellar cell types with observed differences between wild-type and SCA1.**

**Figure S1.1: Purkinje cells (PCs) at 12 and 24 weeks of age.** (a) UMAP of subpopulations of WT and SCA1 mice, and overlay of conditions of mouse Purkinje cells (PC) at 12 weeks. Using a resolution of 0.6, we found 3 subpopulations, whereby population 1 is mostly WT, and 2 and 3 are considered mixed. (b) Mito-processes that are enriched in subpopulations of WT and SCA1 PCs at 12 weeks. Mitochondrial dynamics, OXPHOS (mt), OXPHOS and Ca2+ signaling and transport are the top 4 processes that differ between conditions, whereby condition 3 shows the highest number of implicated genes. (c) Enriched mito-processes based on differentially expressed genes in pseudo-bulk, comparing SCA1 to WT PCs. OXPHOS, as well as Glycolysis are the top processes, with a combined score > 1. (d) Glycolysis and transport genes of the 3 PC subpopulations at 12 weeks. *AldoC* is lowest in subpopulation 2, which is rich in SCA1 PCs. The process Glycolysis was chosen as it is enriched in pseudo-bulk, as well as in single-cell analysis at this time-point.

(e) UMAP of subpopulations of WT and SCA1 mice, and overlay of conditions of mouse Purkinje cells (PC) at 24 weeks. Using a resolution of 0.7, we found 4 subpopulations, whereby population 1 is mostly WT, and population 2 consists of mostly SCA1 PCs; 3 and 4 can be considered mixed. (f) Mito-processes that are enriched in subpopulations of WT and SCA1 PCs at 24 weeks. Ca2+ signaling and transport, mito-carrier, or ROS defense are among top processes that differ between conditions. (g) The only enriched process based on differentially expressed genes in pseudo-bulk comparison of SCA1 vs WT PCs is Glycolysis with a combined score of 2.652. (h) Ca2+ signaling and transport genes of the 4 PC subpopulations at 24 weeks. *Mcu* is reduced in population 1 (mostly WT) and induced in population 2 (mostly SCA1) compared to the other 3 subpopulations, while *Itpr1* behaves in an opposite manner (upregulated in population 1 and downregulated in population 2).

**Figure S3.2**: **Deep cerebellar nuclei (DCN) at 5 and 24 weeks of age.** (a) UMAP of subpopulations of WT and SCA1, and overlay of conditions of mouse DCNs at 5 weeks. With a resolution of 0.7, we found 6 well separated clusters. In the condition overlay, we could observe a significantly higher number of WT DCNs compared to SCA1. Cell population 6 is mostly SCA1, while the other 5 subpopulations are dominated by WT cells. (b) mito-process enrichment of DCS subpopulations at 5 weeks of age. Among the top 4 mito-processes are Amino acid metabolism, Transcription (nuclear), Ca2+ signaling and transport, as well as OXPHOS. (c) When comparing pseudo-bulk data of 5 week DCNs, the only enriched mito-process with a combined score > 1 is Ca2+ signaling and transport (combined score of 1.382). *Itpr2* is significantly reduced, while *Slc25a13* and *Mcur1* are significantly induced. (d) Ca2+ signaling and transport genes of DCN subpopulations of 5 week old mice. *Itpr2* is downregulated, as was observed in the pseudo-bulk comparison in subpopulation 6, while *Slc25a13* could not be detected in this small subpopulation predominantly composed of SCA1 cells.

(e) UMAP of mouse DCNs at 24 weeks, identifying on the one hand subpopulations of WT and SCA1, as well as overlaying the two conditions. 3 subpopulations could be found with a resolution of 0.6. Subpopulation 1 is enriched in SCA1 cells, while 2 and 3 are mixed. (f) Among the top mito-processes that differ between the 3 subpopulations are Ca2+ signaling and transport, mitochondrial Carrier and ROS defense. We could not detect any significantly enriched mito-processes when comparing pseudo-bulk data. (g) Ca2+ signaling and transport genes in DCN subpopulations at 24 weeks. *Slc25a13* and *Itpr2* are again downregulated in subpopulation 1, which is dominated by SCA1 DCNs; and Amino acid metabolism in 24 week DCN subpopulations. Among the top downregulated genes in subpopulation 1 are *Agmat*, *Dglucy* and *Bckdhb*. Upregulated in this population is the gene *Maoa*.

**Figure S3.3: Bergmann Glia (BG) at 18 weeks of age.** (a) UMAP of SCA1 and WT Bergmann glia at 18 weeks, and overlay of conditions. with a resolution of 0.5, we found 3 clusters. The distribution of cells seems shifted, with SCA1 being more present in clusters 1 and 2. (b) Among the top enriched mito-processes in the BG subpopulations are Amino acid metabolism, OXPHOS, Translation and Ca2+ signaling and transport. (c) With comparative analysis of pseudo-bulk data of 18 week BG, we found several mito-processes enriched with a combined score > 1. Among those processes are Glycolysis, Ca2+ signaling and transport, FA degradation and beta-oxidation, Lipid metabolism and TCA cycle. In

Glycolysis, the genes AldoC and Gpi1 are significantly differentially expressed, with *AldoC*, *Slc2a3*, *Eno2* and *Hk2* being down- and *Gpi1* being upregulated. This is similar to the Purkinje cells at this time-point. In Ca2+ signaling and transport, *Itpr1* is downregulated in *SCA1*, while *Itpr2* and *Efhd1* are upregulated, again reflecting the situation of PCs at this age. FA degradation and beta-oxidation shows many significantly upregulated genes and *Acat3*, which is downregulated in SCA1 BGs. The Lipid metabolism genes *Acss3* and *Gpam* are upregulated, while *Prelid3a* is significantly, but weakly reduced. Finally, *Idh1* is downregulated in SCA1 BGs from the mito-process TCA cycle, while *Cs*, *Ogdh* and *Suclg2* are upregulated. (d) For comparative analysis of subpopulations, we decided to show Glycolysis, Ca2+ signaling and transport, as well as Amino acid metabolism. Due to the heterogeneity of the subpopulations, the data are more difficult to analyze. However, we put most emphasis on subpopulation 3, which is predominated by WT cells compared to subpopulations 1 and 2. In Glycolysis, several genes are upregulated in subpopulation 3 compared to the other 2, including *Slc2a1*, *Pgk1* or *Pkm*. *AldoC* is not significantly deregulated, arguing for our observation of highly mixed cell populations in the 3 BG subpopulations. The Ca2+ signaling and transport gene *Itpr1* is strongly and significantly induced in subpopulation 3, while *Efhd1* is induced in subpopulation 1, which is the one with the highest number of SCA1 BGs. Finally, in Amino acid metabolism, the most distinguishing genes can be found in subpopulation 2, including *Chdh, Gls2*, *Ckmt1*, *Aldh18a1*, *Gls*, *Dglucy* and *Pcca*, which all are significantly induced in this subpopulation of 18 week old BGs.

**Supplementary Figure S4.1:**

#### Analysis of subpopulation differences between selected human cerebellar cell types

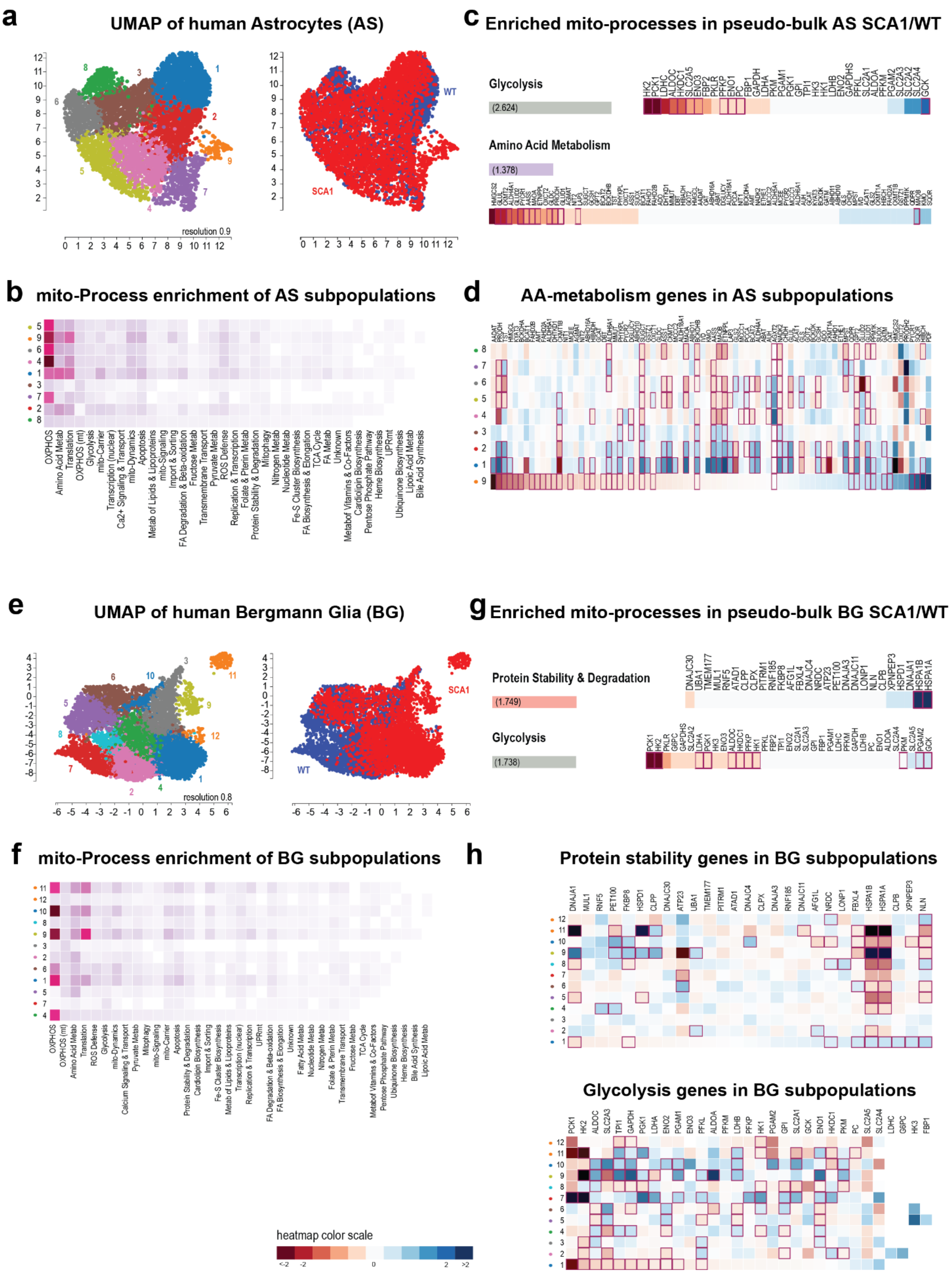

#### Supplementary Figure S4.2:

### Analysis of subpopulation differences between selected human cerrebellar cell types

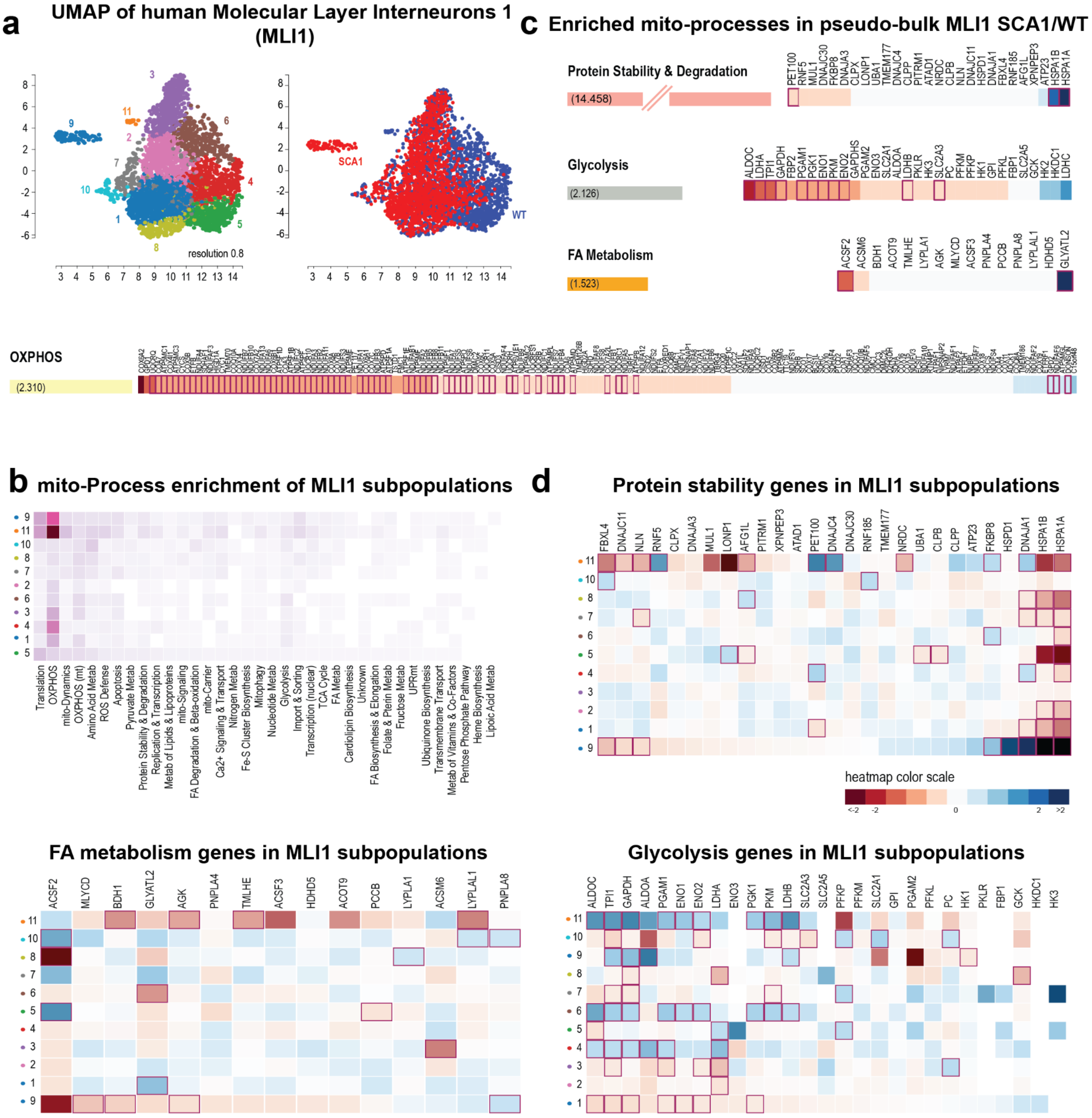

#### Supplementary Figure S4.3:

### Analysis of subpopulation differences between selected human cerrebellar cell types

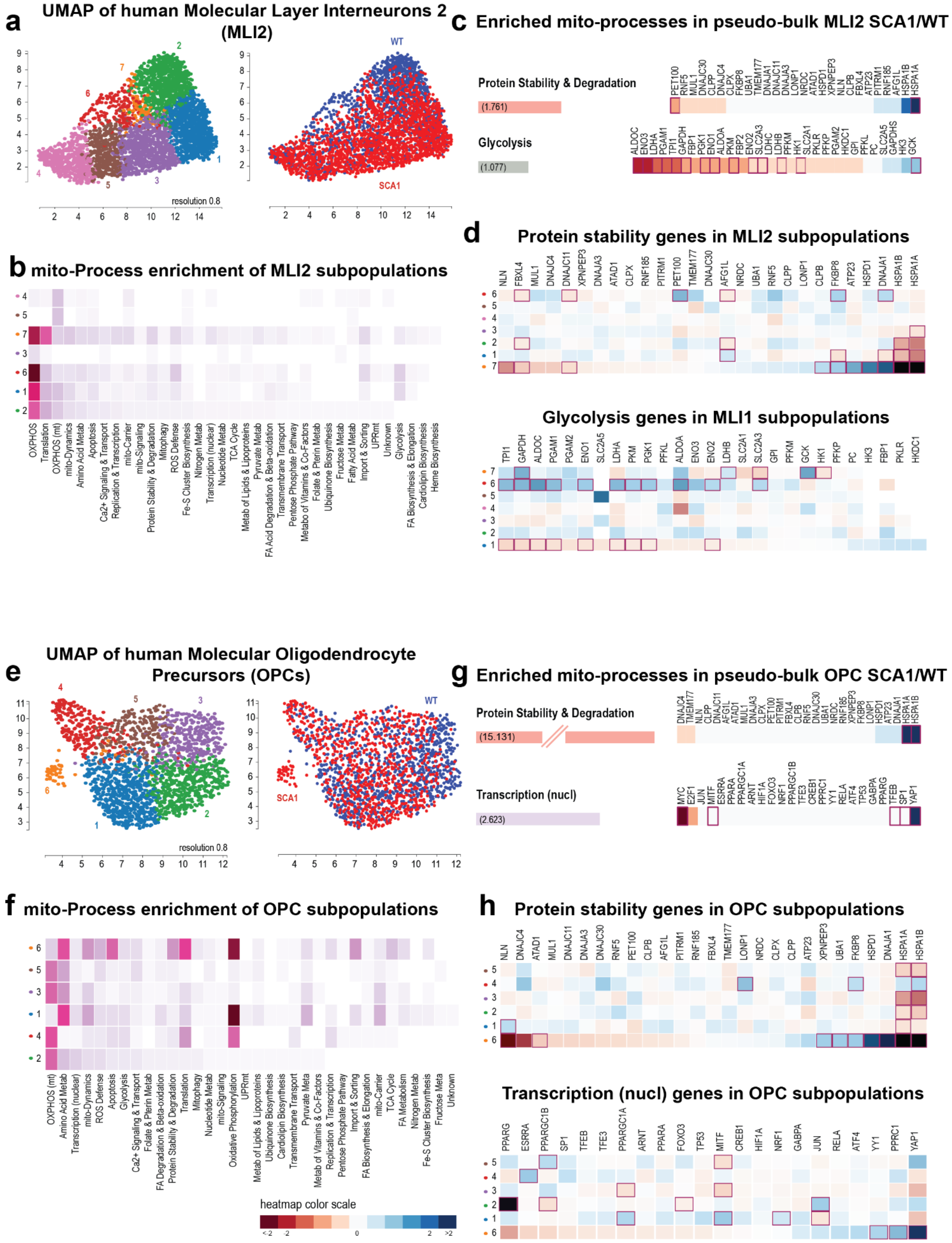

#### Supplementary Figure S4.4:

##### Analysis of subpopulation differences between selected human cerrebellar cell types

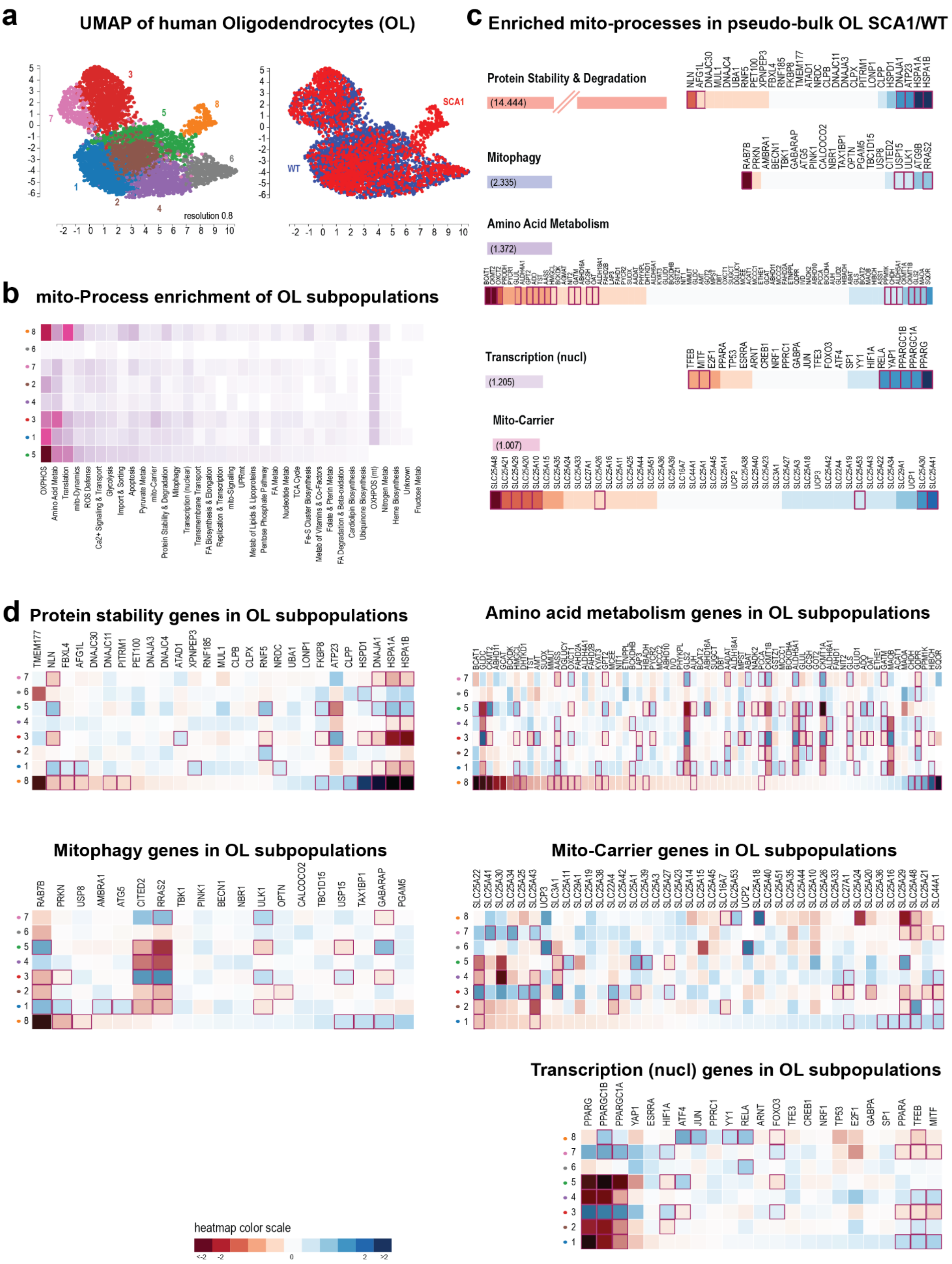

**Supplementary Figure S4: Single-cell and bulk analysis of human cerebellar cell types with observed differences between wild-type and SCA1.**

**Supplementary Figure S4.1: Human Astrocytes (AS) and Bergmann Glia (BG).** (a) UMAP of Astrocytes AS subtypes and overlaid conditions. With a resolution of 0.9, we found 9 subpopulations of AS. Subpopulation 9 is entirely composed of SCA1 AS. (b) Enriched mito-processes in AS subpopulations. Among the most present mito-processes are OXPHOS, Amino acid metabolism, Translation and Glycolysis. (c) The enriched mito-processes Glycolysis and Amino acid metabolism have a combined score > 1 in pseudo-bulk analysis of human AS, comparing SCA1 to healthy control. Among the genes significantly downregulated in Glycolysis are *HK2*, *PCK1*, *ALDOC*, *HKDC1*, but also *ENO3* and *ENO1*. *GCK* is upregulated in SCA1 AS. In Amino acid metabolism, the genes *GLUL*, *CKMT2*, *ALDH4A1*, *PYCR1*, *AASS*, *MAOA*, *ETNPPL*, *GLDC*, *PRODH* and *GLUD1* are significantly reduced in SCA1 AS. *MOAB* on the other hand is upregulated. (d) We chose Amino acid metabolism as the mito-process for single-cell based analysis of subpopulations and sorted according to subpopulation 9, which is composed of SCA1 cells. Among the genes that are significantly reduced in SCA1 cells is the gene PRODH, which was also found in the pseudo-bulk analysis.

(e) UMAP of Bergmann Glia (BG) subtypes and overlaid conditions. With a resolution of 0.8, we could find 12 subclusters. Subpopulations 7 and 8 are predominantly healthy control, while subpopulations 1, 3, 9, 11 and 12 are predominantly SCA1. The other subpopulations are mixed. (f) Among the most enriched mito-processes between the BG subpopulations were OXPHOS, Amino acid metabolism, Translation ROS defense and Glycolysis. (g) When analyzing the pseudo-bulk data of human AS, we found the mito-processes Protein stability and degradation, as well as Glycolysis enriched. The protein stability genes *HSPA1A* and *HSPA1B* showed a very strong upregulation in SCA1 patients. In Glycolysis, the genes *PCK1* and *HK2* showed strong downregulation, as did *PGK1*, *LDHA*, *ALDOC*, *HKDC1* and *HK1*, while *GCK* and *PGAM2* were slightly induced. (h) The same processes, Protein stability and degradation as well as Glycolysi, are shown for single-cell based analysis of BG subpopulations. The chaperons *HSPA1A* and *HSPA1B* were very strongly and also significantly induced in subpopulations 9 and 11, which are both composed of SCA1 cells. In Glycolysis, *PCK1* is strongly reduced in populations 11 (SCA1 BGs) and 1 (mostly SCA1 BGs), while *HK2* demonstrates a more mixed image. *ALDOC* is less affected in these cells.

**Supplementary Figure S4.2: Human Molecular Layer Interneurons 1 (MLI1).** (a) Mito-gene based UMAP of SCA1 and WT MLI1 and overlay of conditions. With a resolution of 0.8, 11 subclusters were identified. There is visible separation between WT and SCA1 cells, with cluster 9 being only composed of SCA1 cells, and clusters 5, 4 and 6 being predominantly WT cells. (b) Mito-processes with significantly differentially expressed genes between the subpopulations. Among the most affected processes are Translation, OXPHOS, mito-Dynamics and Amino acid metabolism. (c) Enriched mito-processes emanating from pseudo-bulk analysis of MLI1 SCA1 vs WT cells. The most enriched mito-process was Protein stability and degradation, with the *HSPA1A* and *HSPA1B* genes highly upregulated in the SCA1 condition, and *PET100* downregulated compared to healthy control. Other processes with combined scores > 1 were Glycolysis, Fatty acid metabolism and OXPHOS. In Glycolysis, *ALDOC*, together with 11 other genes, were significantly reduced in SCA1. Among them were *DHA*, *PGAM1*, *TPI1*, *GAPDH*, *PGK1*, *ENO1*, *PKM* and *ENO2*. In FA metabolism, finally, *ACSF2* is reduced in SCA1 compared to WT, while *GLYATL2* behaves in an opposite way. In OXPHOS, many nuclear-encoded genes involved iN OXPHOS were significantly downregulated in SCA1 MLI1 cells, compared to the healthy individuals. (d) Based on enrichment analysis of the pseudo-bulk data, we looked closer at Protein stability and degradation, FA metabolism and Glycolysis in MLI1 subpopulations. In Protein stability and degradation, the chaperones *HSPA1A* and *HSPA1B* are again strongly upregulated in the SCA1 subpopulation (9, blue dots), as are the *DNAJA1* and *HSPD1* genes. In FA metabolism, *ACSF2* shows the strongest differential expression, though it is non-significantly downregulated in subpopulation 9 (SCA1 cells). In Glycolysis, many genes that have been identified in pseudo-bulk data comparison show also differential expression in MLI1 subpopulations. Among those is *ALDOC*, *GAPDH*, *PGAM1*, *PKM*, or *TPI1*.

**Supplementary Figure S4.3: Human Molecular Layer Interneurons 2 (MLI2) and Oligodendrocyte precursors (OPCs)**. (a) UMAP of Molecular Layer Interneurons 2 subpopulations of SCA1 and healthy control, and overlay of conditions. Using a resolution of 0.8, 7 subpopulations could be identified. Separation of populations is not as clear as with other cell types, though subpopulation 2 and also 6 are mainly composed of healthy control cells and subpopulations 7 of SCA1 cells. (b) Mito-process enrichments of MLI2 subpopulations. Among the top enriched processes are OXPHOS, Translation, mito-Dynamics and Amino amino acid metabolism. (c) When performing enrichment analysis on the pseudo-bulk comparisons, Protein stability and degradation, as well as Glycolysis were identified with a combined score > 1. Again, the chaperones HSPA1A and HSPA1B were strongly enriched in the SCA1 MLI2 cells, while PET100 was reduced. In Glycolysis, a number of genes were downregulated. Among those were, as with MLI1 cells, *ALDOC*, *LDHA*, *PGAM1*, *TPI1*, *GAPDH*, *PGK1*, *ENO1, PKM* and *ENO2*. (d) We looked at the same processes in MLI2 subpopulations and found the *HSPA1A* and *HSPA1B* genes highest in subpopulation 7, together with the *DNAJA1* and *HSPD1* genes. Many Glycolysis genes are upregulated in subpopulations 6 (more cells from healthy control) as compared to for instance subpopulation 1, which is mixed.

(e) UMAP of Oligodendrocyte precursor (OPC) subpopulations of SCA1 and healthy control, and overlay of conditions. 6 subpopulations could be found with a resolution of 0.8, whereby subpopulation 6 is separate from the others and composed of SCA1 OPCs. (f) Mito-processes associated with differentially expressed genes in OPC subpopulations. The top processes include Amino acid metabolism, Transcription (nucl), mito-Dynamics, ROS defense and Apoptosis. (g) Based on the enrichment analyses, we further analyzed Protein stability and degradation, as well as Transcription (nuclear) in OPC subpopulations. The chaperones *HSPA1A* and *HSPA1B* were again induced in the SCA1-only subpopulation (6), together with the *DNAJA1* and *HSPD1* genes. The gene *NLN* (coding for the mitochondrial Oligopeptidase Neurolysin) and *ATAD1* on the other hand were reduced in this subpopulation compared to the others. In the nuclear transcriptional regulation process, subpopulation 1 showed significant induction of the *YAP1* transcription factor, together with the *YY1* and *PPRC1* genes.

**Supplementary Figure S4.4: Human Oligodendrocytes (OL).** (a) UMAP of Oligodendrocyte (OL) subpopulations of SCA1 and healthy control, and overlay of conditions. We found 8 subpopulations with a resolution of 0.8. Subpopulation 8 is composed of SCA1 OLs, while the other populations can be considered mixed. (b) We found several mito-processes that could be associated with subpopulations of OLs. Among the top ones are OXPHOS, Amino acid metabolism, Translation, mitochondrial Dynamics, ROS defense or Ca2+ signaling and transport. (c) Enriched mito-processes of pseudo-bulk comparison between the two conditions included Protein stability and degradation, Mitophagy, Amino acid metabolism, Transcription (nuclear) and mitochondrial Carrier with a combined score > 1. The chaperones *HSPA1A* and *HSPA1B* are again strongly induced in SCA1 cells, as is the DNAJA1 gene. The mitophagy gene *RRAS2* is induced, while *RAB7B* is strongly downregulated in the mutation situation. Quite a few genes involved in Amino acid metabolism are reduced in SCA1, including *BCAT1*, *CMKT2*, *OXCT2*, *GLUL*, *GPT2*, *ADO*, *TST* and *AASS*. Among the upregulated genes of this process are *MAOA*, *GLS2* or *CKMT1B*. The nuclear transcriptional regulator gene *PPARG* together with *PPARGC1A*, *PPARGC1B*, *YAP1* and *RELA* are induced in SCA1 OLs, while *TFEB* and *MITF* are downregulated. Finally, in the process mitochondrial Carrier, the *SLC25A48*, *SCL25A21*, *SCA25A29*, *SLC25A20* and *SLC25A10* genes are reduced, while *SLC25A30* and *SLC25A41* are upregulated in the mutant OLs. (d) We looked at the same processes in OL subpopluations. *HSPA1A* and *HSPA1B* were together with *DNAJA1* and *HSPD1* again strongly induced in the SCA1 subpopulation 8 in the Protein stability process. In Mitophagy, the *RAB7B* gene was strongly reduced in the same subpopulation. We also found strong overlap between the pseudo-bulk and single-cell analysis, focusing on the subpopulation 8 composed of SCA1 cells. Among the downregulated genes are the two genes *BCAT1* and *CKMT2*. In the mitochondrial Carrier process, the two genes *SLC25A29* and *SLC25A48* are strongly reduced in the SCA1 only subpopulation. Finally, in the process nuclear Transcription, we find the genes *PPARGC1B*, *YY1* and *RELA* induced, and *TFEB* reduced in subpopulation 8 that is composed of SCA1 cells.

**Supplementary Figure S5:**

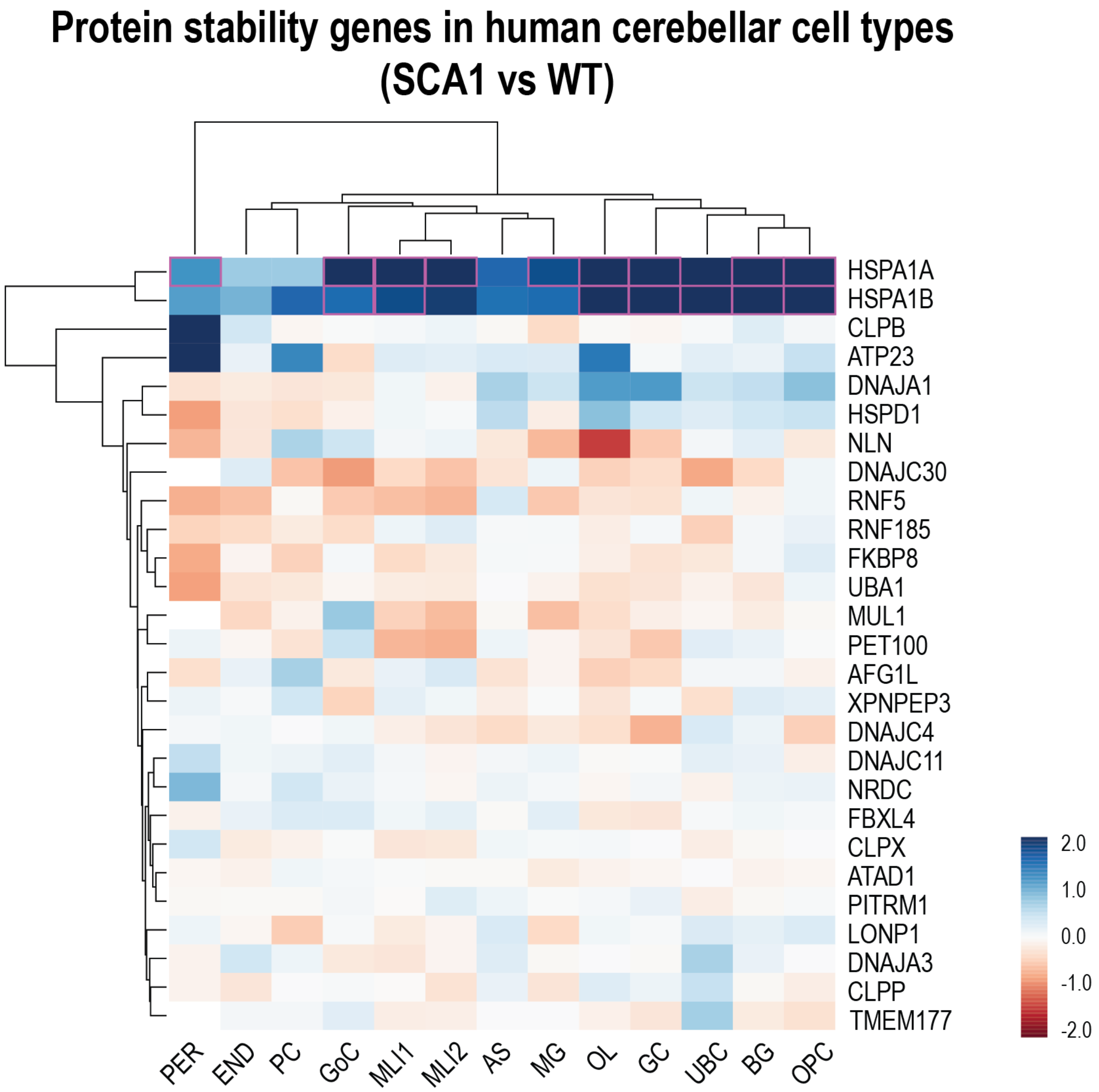


**Supplementary Figure S5:** Heatmap of the mito-process Protein stability and degradation over all human cerebellar cell types. The two genes *HSPA1A* and *HSPA1B* are strongly and in most cell types also significantly induced in SCA1 compared to WT.

#### Supplementary Figure S6: Usage instructions for mitoXplorer 3.0 data mining

1. mitoXplorer 3.0 single-cell data mining of available datasets starts by going to the 'database' menu item (a), selecting 'Single Cell Data' (b), selecting the Organism (c) and the Project (d).

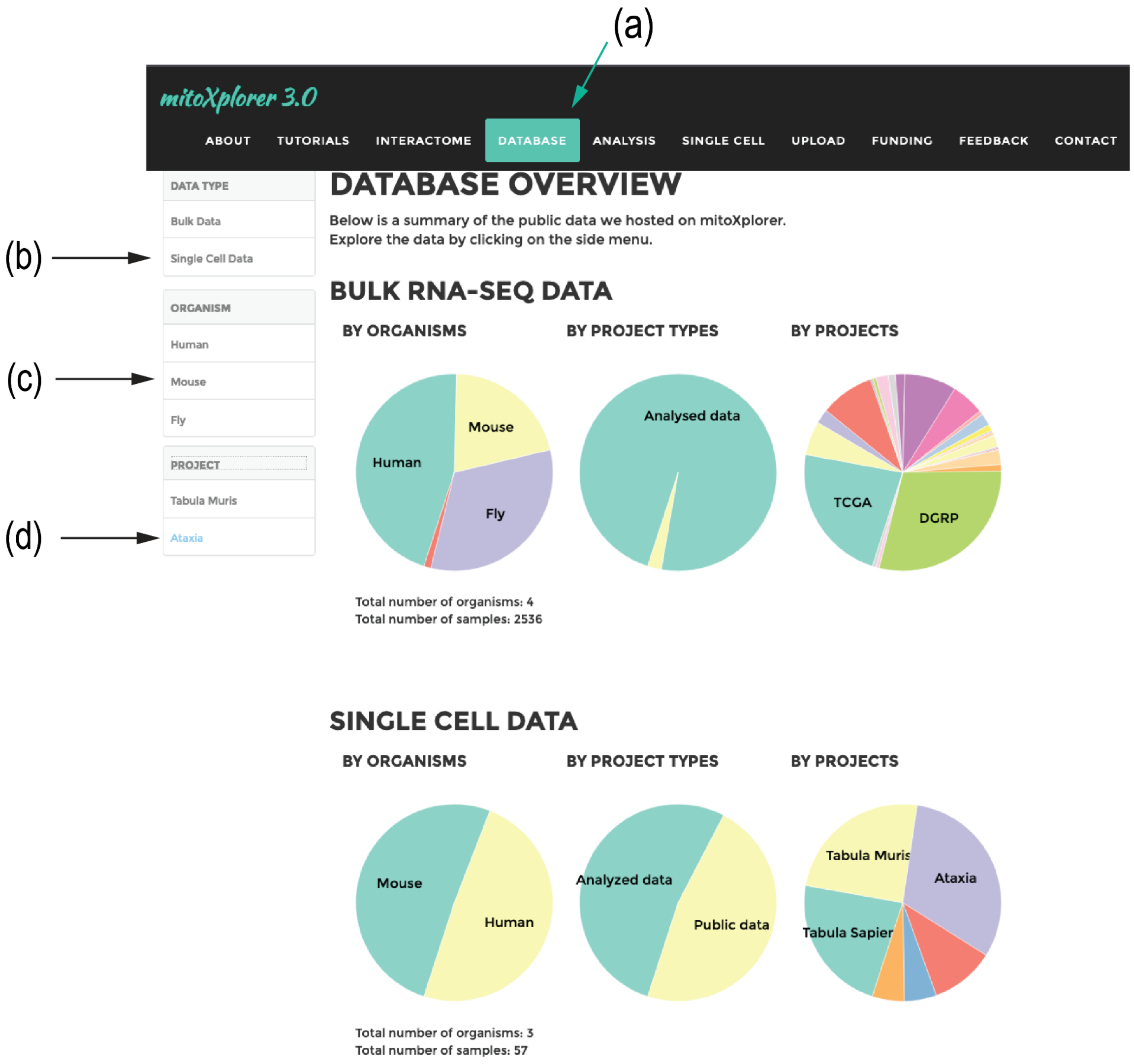

**2** This will open up a page with tabular information describing the datasets available for the chosen project. Now choose the dataset you want to mine (a).

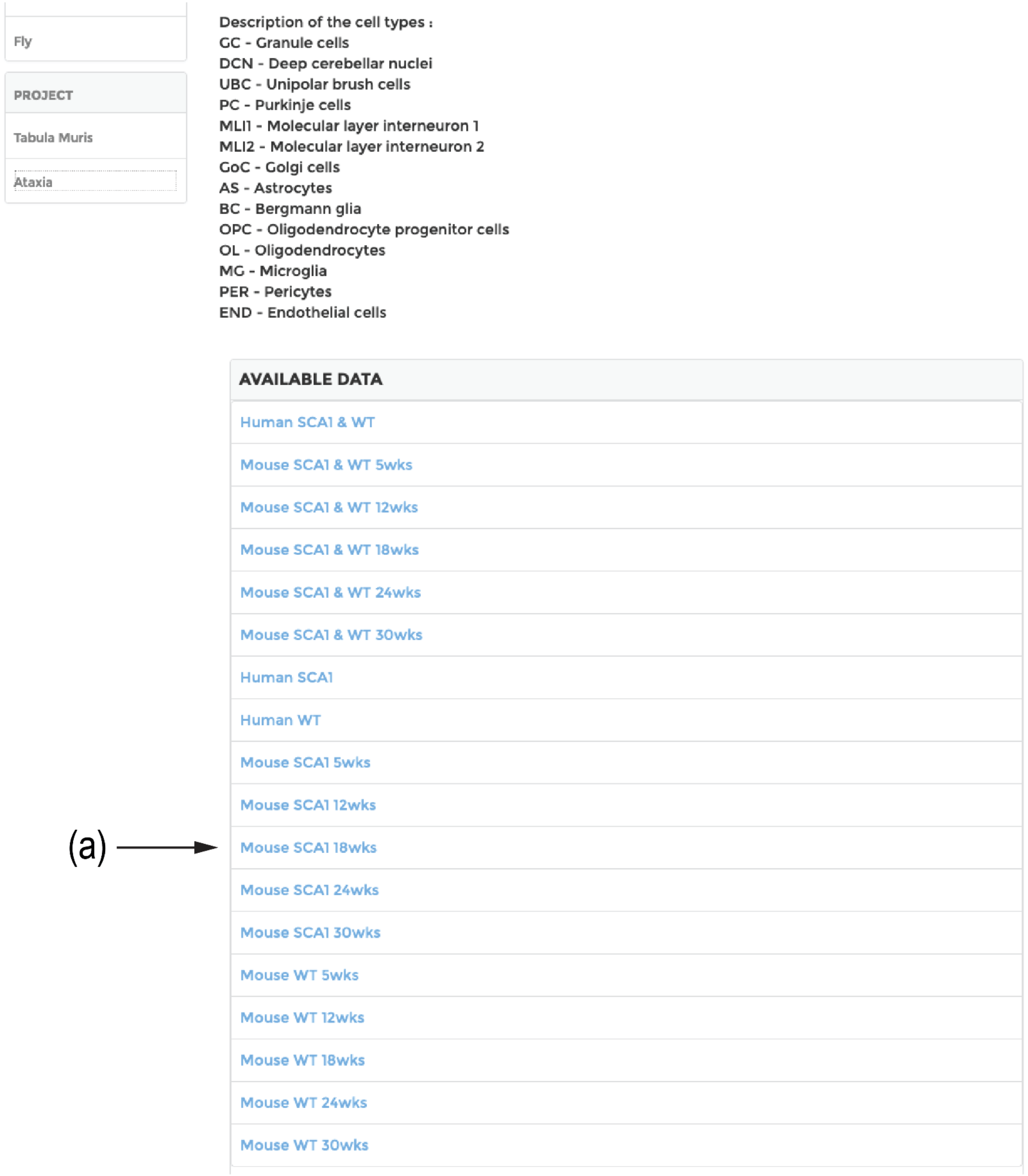

**3** The original UMAP will appear that provides the distribution of cell types in the sequenced dataset. From this point onwards, steps are similar between provided and uploaded datasets. Select here the cell type (a) and the clustering resolution (b) and press cluster (c) to perform subpopulation clustering.

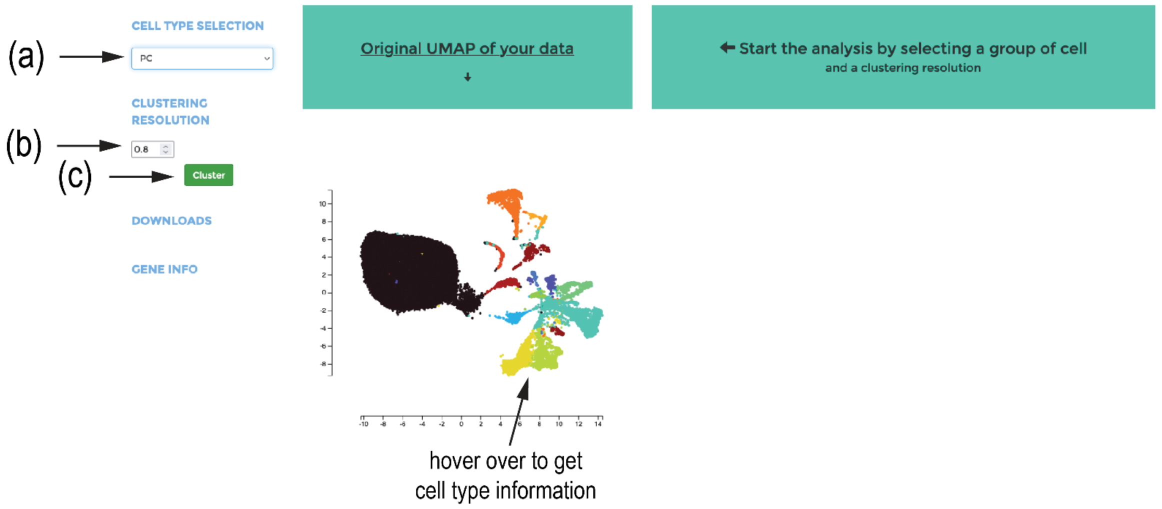


**4** Here the mito-gene specific UMAP will be displayed next to the original UMAP. Hovering over the colored dots will reveal the subpopulation number. In case of using 2 conditions, select the radio buttion 'conditions' to display the overlay between the conditions.

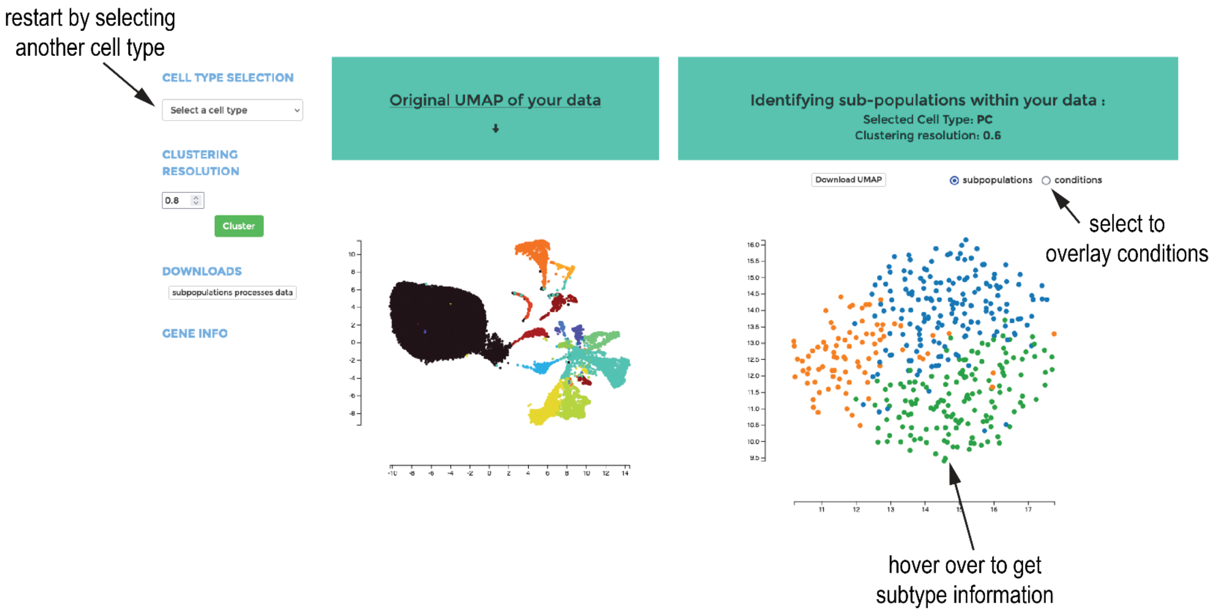

**5** Below the UMAP after cell type selection and mito-gene based subclustering, a stacked bar plot will appear that contains the mito-processes associated with the cell types. the size of the colored stack represents the number of significant genes in that process in the cell subtype. Press on one of the horizontal bars to display the selected process for gene-based analysis with a heatmap (point 6).

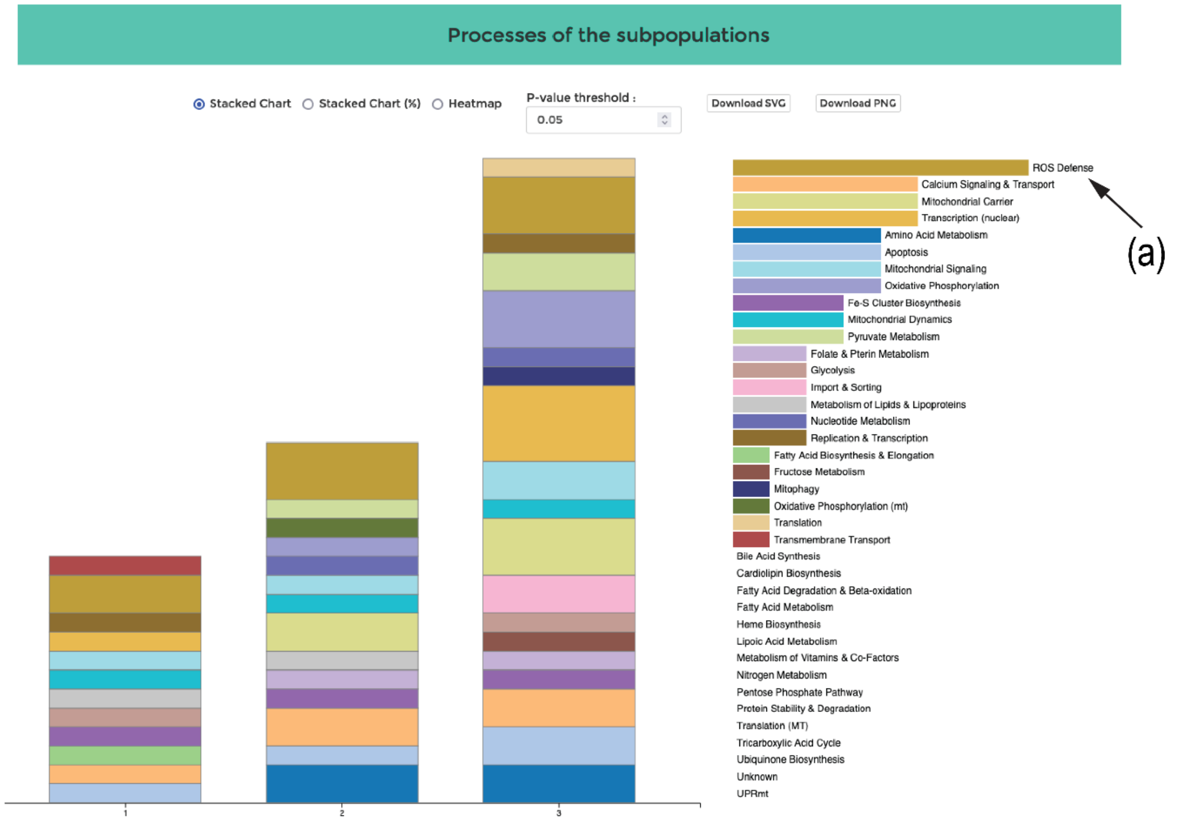


**6** An interactive sortable heatmap shows the individual genes in a selected mito-process, here ROS defense, for the cell subtypes. Click on one of the subtype numbers to resort the heatmap according to the expression levels of the genes in this suptype. Hover over one of the bars to get information on raw & normalized read counts, log2FC and p-value.

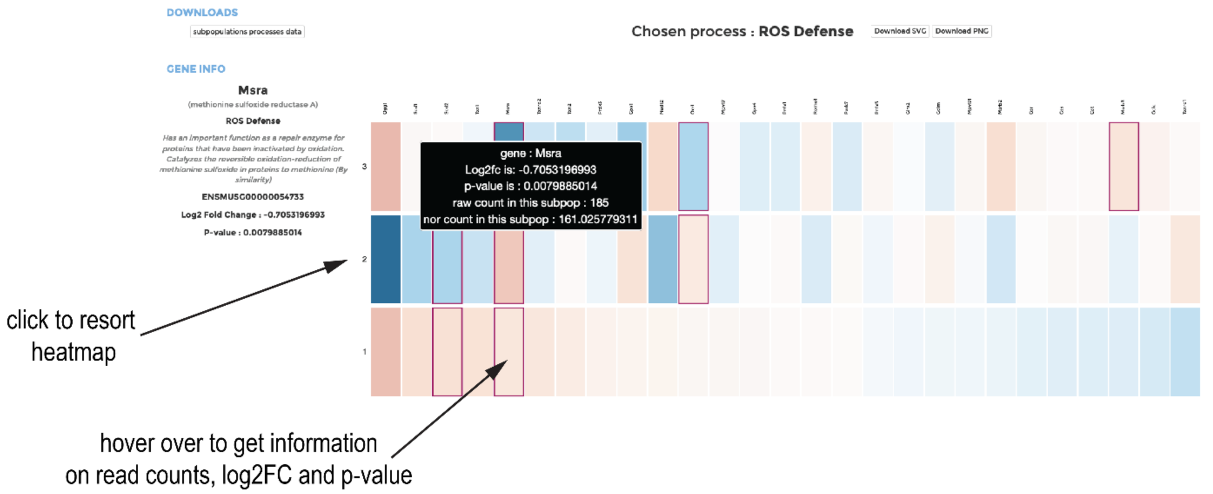

7 When choosing the stacked chart (%) (a), the mito-process to be viewed must be selected by a drop-down menu (b).

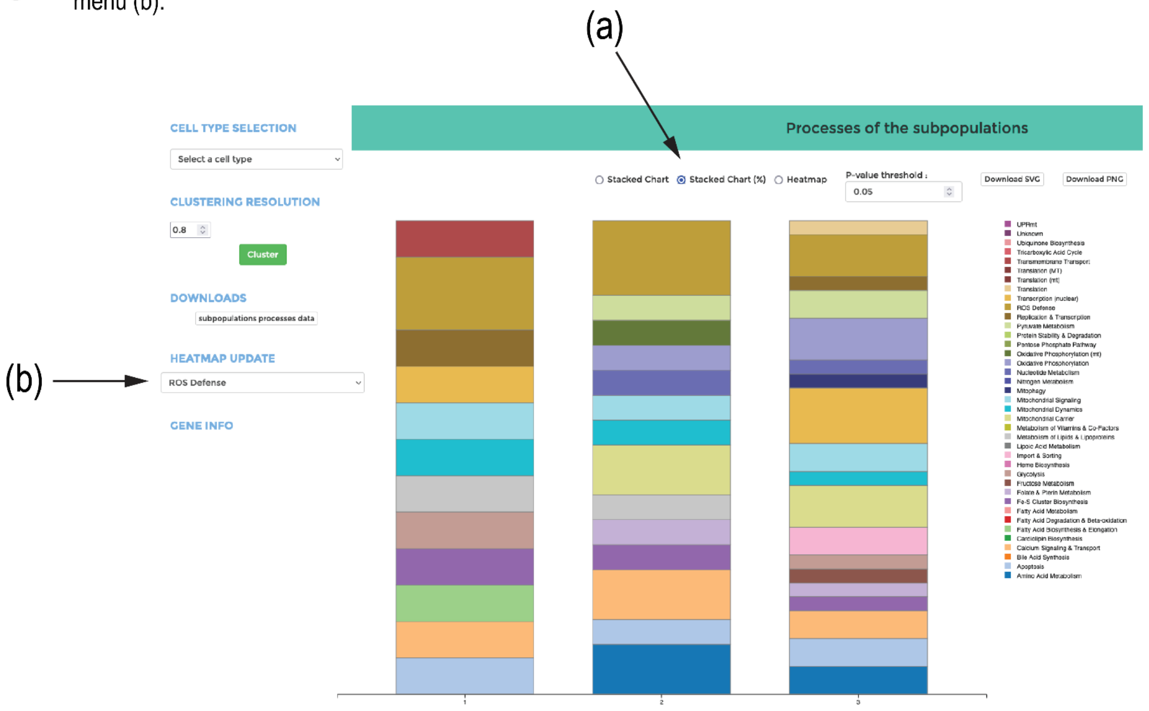


8 The same applies to the 'Heatmap' display (a), where the mito-process must be chosen by the drop-down menu (b).

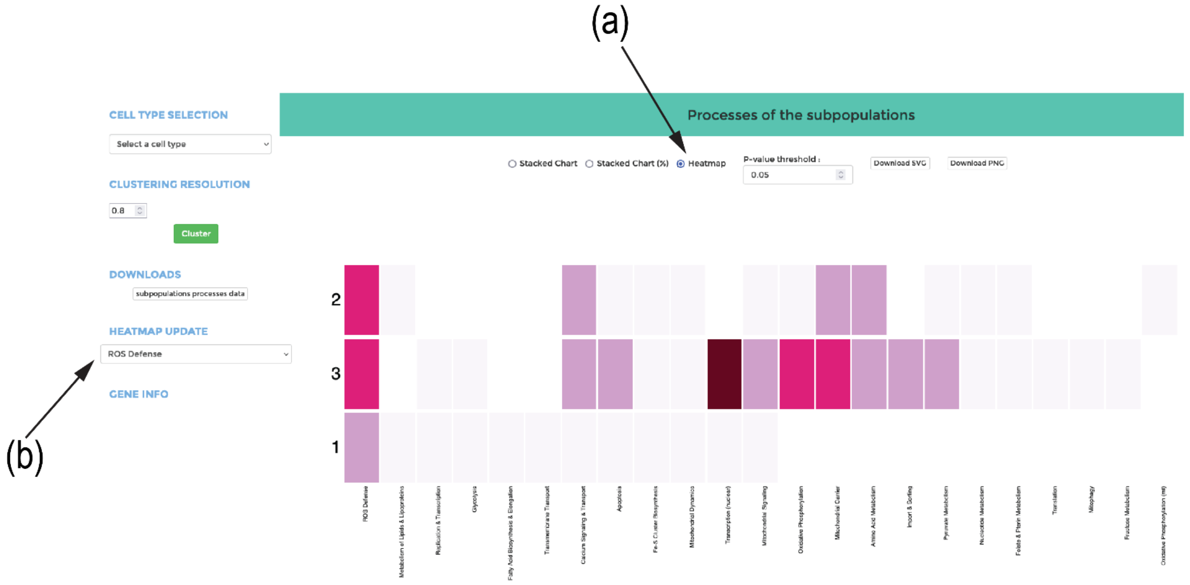
